# Region-specific generation and retention of acetylated cohesin shape the genome-wide pattern of sister chromatid cohesion

**DOI:** 10.64898/2026.01.14.699591

**Authors:** Mai I. Shimizu, Masashige Bando, Atsunori Yoshimura, Toyonori Sakata, Tomoko Nishiyama, Katsuhiko Shirahige, Takashi Sutani

## Abstract

Sister chromatid cohesion depends on ESCO2-mediated acetylation of the cohesin subunit SMC3, yet how this modification is established and maintained across the genome remains poorly understood. Here, we examined the chromatin-binding dynamics of ESCO2 and its enzymatic activity using quantitative ChIP-seq of human cells synchronously progressing through S phase. ESCO2 bound broadly along chromosomes without forming discrete peaks and preferentially localized to transcriptionally inactive regions, but was not confined to H3K9me3-or H3K27me3-enriched heterochromatin. Consistent with this binding pattern, ESCO2-dependent SMC3 acetylation showed a similarly broad distribution in these regions. In contrast, ESCO1, a paralog of ESCO2, primarily mediated SMC3 acetylation at cohesin peak sites. Notably, most of the broadly distributed, ESCO2-dependent SMC3 acetylation generated during S phase was progressively lost from chromatin after replication, whereas the fraction that persisted into late G2 accumulated in domains enriched for the repressive H3K27me3 modification. Our work identifies selective post-replicative retention of acetylated cohesin as a previously unappreciated step in shaping the genome-wide architecture of sister chromatid cohesion.

## Introduction

In proliferating eukaryotic cells, sister chromatids that arise from DNA replication remain physically tethered until mitotic anaphase, and this sister chromatid cohesion is indispensable for faithful chromosome segregation and genome stability (1). Cohesin, a ring-shaped protein complex, mediates this process by topologically entrapping the two chromatids (2–5). Although cohesin also regulates higher-order chromosome organization and gene expression during interphase (2, 6), the subset of cohesin complexes that confer sister chromatid cohesion possesses a defining molecular feature: acetylation of the SMC3 subunit on conserved lysines, corresponding to K105 and K106 in human SMC3 (1, 7–9). This modification is catalyzed by evolutionarily conserved acetyltransferases, Eco1 in yeast and ESCO1/ESCO2 in human cells, and its establishment in a DNA replication-coupled manner is thought to be essential for converting dynamically associating cohesin into a stably chromatin-bound, cohesion-mediating form (1, 9). SMC3 acetylation antagonizes cohesin’s interactions with the loader NIPBL and the releaser WAPL (10–13), and in human cells it further promotes binding of the cohesion-stabilizing factor Sororin (14). As a result, cells impaired in SMC3 acetylation fail to establish stable cohesion and undergo chromosome segregation defects and catastrophic mitoses (7, 8, 15).

The two human SMC3 acetyltransferases, ESCO1 and ESCO2, exhibit markedly distinct intracellular behaviors. ESCO1 is expressed throughout the cell cycle, whereas ESCO2 is restricted to S phase and is degraded once DNA replication is complete (16, 17). Previously, we demonstrated that ESCO1 localizes to cohesin binding sites (e.g. CTCF-anchored sites) in the genome throughout interphase via an interaction with PDS5A or PDS5B, enabling ESCO1 to acetylate cohesin even outside of S phase (18). In contrast, ESCO2’s activity is independent of PDS5 (18) and tightly linked to DNA replication, like yeast Eco1. Studies in *Xenopus* showed that the prereplication complex recruits XEco2 (the *Xenopus* ESCO2 ortholog) to chromatin as DNA replication initiates (19), and works in human cells revealed ESCO2 is loaded at replication sites through the interaction with the MCM replicative helicase (17, 20) and the replication factor PCNA (21). These observations suggest that ESCO2 travels with replication forks to acetylate cohesin complexes in their vicinity.

Beyond these differences in regulation, ESCO1 and ESCO2 contribute to sister chromatid cohesion at distinct chromosomal regions. In cells lacking both ESCO1 and ESCO2, sister chromatid cohesion is completely lost (16, 18). In contrast, human cultured cells deficient in either ESCO1 or ESCO2 alone exhibit distinct alterations in mitotic chromosome morphology.

Specifically, loss of ESCO1 leads to the separation of chromosome arms, whereas loss of ESCO2 causes defects in cohesion near centromeric regions (20). The notion that ESCO2 strongly contributes to cohesion at the centromeric heterochromatin is supported by observations from patient cells with *ESCO2* mutations (Roberts syndrome), *ESCO2* knockout mice, and chicken DT40 *ESCO2*-knockout cells (22–24). These findings indicate that ESCO1 and ESCO2 differ not only in their regulation but also in the chromosomal regions where they function. However, in standard ChIP-seq analyses, ESCO2 exhibits only a few clear, reproducible binding peaks (20, 25), leaving it unresolved whether its localization is associated with heterochromatin.

Recent advances in chromosome-conformation capture and imaging methods have challenged the classical view that cohesion is uniformly distributed along replicated chromosomes. Sister-chromatid-sensitive Hi-C (scsHi-C) (26) enables sister chromatid contacts to be mapped at high resolution on a genome-wide scale and revealed that trans-sister contacts are preferentially enriched near the boundaries of topologically associating domains (TADs), such as CTCF-anchored sites. Complementary super-resolution imaging further demonstrated that sister chromatids are frequently paired at discrete genomic positions in a cohesin-dependent manner (27). A notable exception, revealed by scsHi-C, is the presence of strong intra-TAD sister contacts within TADs enriched for H3K27me3 (26), indicating that facultative heterochromatin forms cohesive structures that differ markedly from those in euchromatic domains. Together, these findings suggest that cohesion is established in a highly nonuniform, chromatin-state-dependent manner. How such a genomic distribution of cohesion is generated, and how ESCO1 and ESCO2 contribute differentially to these distinct cohesion patterns, remains unresolved.

In this study, we investigated the chromatin-binding dynamics of ESCO2 and its enzymatic function (i.e., SMC3 acetylation) using quantitative ChIP-seq analysis of human cultured cells synchronously progressing through S phase. We found that ESCO2 binds broadly along chromosomes without forming discrete peaks and preferentially localizes to transcriptionally inactive regions, but is not confined to H3K27me3-rich chromatin. ESCO2-dependent SMC3 acetylation shows a similarly broad distribution in these regions. Notably, most ESCO2-dependent SMC3 acetylation generated during S phase is progressively lost from chromosomes, whereas the SMC3ac that persists into late G2 is enriched in genomic regions marked by high levels of H3K27me3. The region-specific retention of SMC3ac we observed could explain how the genome-wide pattern of sister chromatid cohesion is established. Together, our findings refine the current understanding of how cohesion architecture is spatially and temporally regulated in human cells.

## Materials and Methods

### Reagents and biological resources

For a full list of reagents and biological resources, please see Supplementary Table S1.

### Cell culture and synchronization

Human HeLa S3 and mouse C2C12 cells were cultured in high-glucose D-MEM (Dulbecco’s Modified Eagle’s Medium) supplemented with 10% fetal bovine serum and penicillin-streptomycin-L-glutamine solution. Cell cycle synchronization of HeLa S3 cells was achieved by arresting cells using double thymidine block (16 h in the presence of 2 mM thymidine, 9–10 h without thymidine, and 14–15 h in the presence of 2 mM thymidine), followed by release into thymidine-free medium. The cells were harvested at 0, 3 and 7 h after the second release to prepare cells synchronized to the early S, mid S and G2 phases. To prepare cells arrested at late G2 phase, the Cdk1 inhibitor RO-3306 was added to the culture medium at the final concentration of 10 μM 7 hours after the second release from the thymidine block and incubated for an additional 1.5 hours. For EZH2 inhibition, EPZ-6438 was added to the culture medium at a final concentration of 2 μM 24 h before the first thymidine block and maintained throughout the synchronization procedure until cell harvest.

### Flow cytometry

Cell cycle synchronization was assessed by flow cytometry analysis of DNA content. A portion of the collected cells was fixed by adding ethanol to a final concentration of 70%. The fixed cells were then washed twice with PBS, treated with 50 µg/mL RNase A (Roche), and stained with 50 µg/mL propidium iodide. The DNA content of the stained cells was measured using a BD Accuri™ C6 Flow Cytometer (Becton Dickinson), and the obtained data were analyzed using FlowJo (Becton Dickinson).

### RNA interference

Stealth siRNAs pre-designed and synthesized by Thermo Fisher Scientific were utilized. The sequences of the siRNAs are described in Supplementary Table S1. siRNA transfection was carried out using Lipofectamine RNAiMAX at a final concentration of 50 nM. ESCO1 siRNA was transfected concurrently with the first thymidine treatment, while ESCO2 siRNA was transfected at the time of the first thymidine release. MCM7 siRNA was transfected 8 h prior to the first thymidine treatment. Flow cytometry analysis confirmed that cell cycle progression was unaffected under knockdown conditions.

### Protein analysis

The cells were collected and washed twice with PBS. The cells were then lysed on ice for 10 min in buffer A (25 mM Tris-HCl, pH 7.5, 100 mM NaCl, 5 mM MgCl₂, 10% glycerol, 0.2% NP-40, 1 mM DTT, 10 mM sodium butyrate, cOmplete protease inhibitor cocktail, and PhosSTOP phosphatase inhibitor cocktail) to obtain total cell lysate. Chromatin-enriched fraction was obtained by subjecting the total cell lysate to low-speed centrifugation at 1,500 ×g, followed by washing and resuspending the resulting pellet in buffer A. Both the total cell lysate and chromatin fraction were sonicated using a TOMY UR-21P to shear DNA, followed by boiling in SDS-PAGE sample buffer. Proteins were analyzed by western blotting with Fusion FX (Vilber Bio Imaging) image detection system. Quantitative western blotting was performed using the Jess automated western blotting system (ProteinSimple) according to the manufacturer’s instructions, with acetylated SMC3 (SMC3ac) and SMC3 detected by chemiluminescence and fluorescence, respectively.

### Chromatin immunoprecipitation (ChIP)

ChIP was performed as previously described (28). Briefly, HeLa S3 cells were crosslinked with 1% formaldehyde for 10 min, followed by quenching with 125 mM glycine. The fixed cells were lysed with detergent, chromatin-enriched fraction was obtained by centrifugation, and DNA was sheared by sonication (Sonifier 250D, Branson). Fragmented chromatin was incubated overnight at 4°C with antibody-conjugated protein A or protein G Dynabeads. 5 µg of antibody was used per 10^7^ cells. After this, the beads were washed thoroughly and incubated with elution buffer (50 mM Tris, 10 mM EDTA, and 1% SDS) for 20 min at 65°C to elute DNA. The eluates were incubated at 65°C overnight to reverse crosslinks, and treated with RNase A and subsequently with proteinase K. The samples were purified by QIAquick PCR purification kit according to the manufacturer’s instructions, which yielded ChIP DNA fraction. An aliquot of the chromatin fraction before immunoprecipitation was also subjected to reverse-crosslink, deproteinization, and purification to yield the input DNA fraction. Both ChIP and input DNA were subsequently used for sequencing library preparation (ChIP-seq) or quantitative PCR measurement (ChIP-qPCR). For calibrated (spike-in) ChIP-seq, the mouse-derived chromatin was prepared from asynchronous mouse C2C12 cells using the same cell fixation procedure and mixed with human HeLa cell chromatin prior to chromatin shearing by sonication. The mouse and human chromatin were mixed at a ratio of 1:3 or 1:5, and this ratio was consistently maintained within each series of comparative experiments. Calibrated Sororin ChIP-seq was performed using HeLa cells expressing GFP-tagged mouse Sororin (29). Cells isolated from the bone marrow and spleen of H2B-GFP mice, generated by crossing R26R-H2B-EGFP mice (30) with Vasa-Cre mice, were used as the spike-in control.

### Quantitative PCR (qPCR)

qPCR analysis of ChIP DNA was conducted using a CFX384 Real-Time System (Bio-Rad) or a StepOnePlus Real-Time PCR System (Thermo Fisher Scientific) with the KAPA SYBR FAST qPCR kit, according to the manufacturers’ instructions. Each sample was analyzed in duplicate or triplicate, and the standard deviation was calculated. The primer pairs utilized are listed in Supplementary Table S2.

### ChIP-seq library preparation and sequencing

Both input and ChIP fractions of DNA were processed. Purified DNA was further fragmented to the size of ∼150 bp by the M220 focused-ultrasonicator (Covaris) and subjected to library preparation using the NEBNext Ultra II DNA Library Prep Kit for Illumina. Sequencing of the library was performed using the HiSeq 2500 (Illumina) to produce 65-bp single-end reads or NovaSeq X plus (Illumina) to produce 150-bp paired-end reads.

### Calibrated ChIP-seq data analysis

#### ⁃Mapping

Mapping was performed using Bowtie 2 (v2.3.4.1) (31) with default parameter settings (i.e., allowing no mismatches and reporting the best alignment per read). The human reference genome (UCSC hg19) and the mouse reference genome (UCSC mm10) were used. Reads that mapped uniquely to the human genome but not to the mouse genome were used for the downstream analyses described below. Conversely, reads that mapped only to the mouse genome were used later for normalization purposes. Redundant reads, which mapped to multiple locations in the human or mouse genome, were assumed to originate from repetitive genomic regions and were excluded from downstream analyses. Additionally, reads that mapped to the exact same position in the reference genome were regarded as PCR duplicates and collapsed into a single read. A summary of read mapping statistics is provided in Supplementary Table S3.

#### ⁃Calibration

For a specific ChIP sample, the number of filtered reads mapped to the human and mouse genomes in the ChIP and corresponding input fractions was used to calculate the Occupancy Ratio (OR) according to the following formula, as described previously (32).

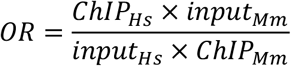

where ChIPHs is the number of reads mapped to the human genome in the ChIP sample; inputHs is the number of reads mapped to human genome in the input sample; ChIPMm is the number of reads mapped to mouse genome in the ChIP sample; and inputMm is the number of reads mapped to mouse genome in the input sample.

The filtered reads mapped to the human genome were binned using parse2wig in the DROMPA package (version 2.5.3) (33), and the result was output as a file in wig format. For an input sample, bin-wise read counts were normalized using the default setting, with the total read count scaled to 20 million. For a ChIP sample, the following normalization option was applied:’-n GR-np X’, where X is the product of 20 million and the corresponding OR. The genome bin size was set to 100 bp, 10 kb, or 100 kb, depending on the downstream analysis.

The computed OR values and normalization parameters are listed in Supplementary Table S3.

#### ⁃ChIP-seq peak calling

ChIP-seq peaks, representing genomic loci with enriched binding of the target protein, were identified with the drompa_peakcall function operated in PC_SHARP mode of DROMPA (33). Read count data binned at 100 bp resolution were used. RAD21 ChIP-seq peaks were called independently for early S, mid S, and G2 samples. Genomic regions that were identified as peaks in all three conditions were defined as RAD21 peak regions and used for downstream analyses in this study.

#### ⁃Normalized fold-enrichment (nFE) calculation

To detect non-localized DNA binding of ESCO2 and MCM7, characterized by broad peak-less ChIP-seq signals, calibrated and binned wig files generated from the corresponding input and ChIP samples of a given ChIP-seq experiment were processed using the PC_ENRICH function in DROMPA. This function generated a file containing the normalized ChIP/Input fold-enrichment (nFE) ratio for each genomic bin, in either bigWig or bedGraph format. Read count data binned at 100 kb resolution were used. To calculate the nFE of non-localized SMC3ac outside cohesin peak regions, we used ChIP and input read count data binned at 100 bp resolution. A read count of zero was assigned to any bin that overlapped, even partially, with RAD21 ChIP-seq peaks. Subsequently, the data were aggregated into 100 kb windows by summing ChIP and input reads within each window, and nFE values were then calculated as the ratio of these sums. ChIP-seq signals resulting from non-localized DNA binding are relatively weak, and the corresponding nFE values contain a substantial amount of background noise derived from non-specifically immunoprecipitated DNA. To facilitate interpretation of the results, the nFE values from a series of comparative experiments were rescaled. For ESCO2 and MCM7, the nFE values were rescaled such that the genome-wide average in the G2 phase equaled 1. For SMC3ac, the nFE values were rescaled so that the average genome-wide signal across early S, mid S, and G2 samples under ESCO1 and ESCO2 double-knockdown equaled 1.

To calculate the nFE of SMC3ac within cohesin peak regions, we used ChIP and input read count data binned at 100 bp resolution. Bins that overlapped, even partially, with RAD21 ChIP-seq peaks were extracted, and the total number of ChIP and input reads within these bins was summed separately. The nFE value was then calculated as the ratio of these totals.

#### ⁃Visualization

The distribution of ChIP read counts or ChIP/input ratio across the genome was visualized using the Integrative Genomics Viewer (34) and the drompa_draw function of DROMPA. In DROMPA, the PC_SHARP and PC_ENRICH modes were used for the respective visualizations. Heatmaps were generated using the computeMatrix and plotHeatmap functions in deepTools (35), based on the output generated using 100 bp binning.

### Calculation of the fraction replicated

In ChIP-seq experiments using cell samples synchronized at specific cell cycle phases, we defined the fraction replicated (% replicated) of a given genomic region as the proportion of cells in which that genomic region has been replicated. This metric was calculated using a previously reported method (36), in which the DNA content at each genomic locus is assessed relative to a reference sample. In this study, we used input sequencing data from ChIP-seq experiments to estimate DNA content, and cells arrested in late G2 phase with the Cdk1 inhibitor RO-3306 served as the reference. Read counts were binned at a resolution of 100 kb using the parse2wig function in DROMPA, and the DNA content ratio for each bin was calculated by comparing the sample of interest to the late G2-arrested reference. Subsequently, the calculated ratios were transformed into % replicated values using the following equation.

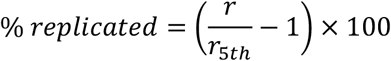

where r is the DNA content ratio at the given genomic bin; and r5th is the 5^th^ percentile of the DNA content ratio in the genome. For samples collected 7 hours after thymidine release (corresponding to G2 phase), in which DNA replication is completed genome-wide, the calculated ratios were transformed into % replicated values using a different equation.

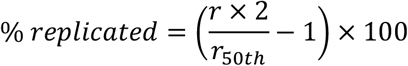

where r is the DNA content ratio at the given genome bin; and r50th is the median of the DNA content ratio in the genome.

Based on % replicated values in early-S and mid-S cells, genomic regions were classified into three categories: ERR (early replicating regions), MRR (mid replicating regions), and LRR (late replicating regions). Regions with % replicated values exceeding 20% in early S phase were defined as ERR. Among the remaining regions, those with % replicated values exceeding 20% in mid S phase were defined as MRR. All other regions were classified as LRR.

Replication timing was evaluated using the ENCODE wavelet-smoothed Repli-seq signal, a continuous replication timing score derived from weighted averaging of six Repli-seq fractions followed by wavelet smoothing (37).

### Experimental Replication

ESCO2 ChIP-seq in mid S, MCM7 ChIP-seq in early S, and SMC3ac ChIP-seq in late G2 were performed in two independent biological experiments. After confirming a reproducible overall distribution between the replicates, one of the datasets was used for detailed downstream analysis and data visualization. Other ChIP-seq conditions were analyzed in one biological experiment and were supported, where indicated, by ChIP-qPCR or biochemical analysis.

### Data visualization and statistical analysis

Data visualizations, including scatter plots, boxplots, aggregation plots, and correlation matrices, were generated using R (38) and Microsoft Excel. The ggplot2 and corrplot packages were used for visualization in R. For clarity, outliers were excluded from some scatter plots; in all cases, these accounted for less than 5% of the total data points. Statistical significance was assessed using two-sided Mann–Whitney *U* tests in R.

### Publicly available sequencing data

Publicly available ChIP-seq and RNA-seq datasets for HeLa S3 cells were downloaded from public databases and used in this study. ChIP-seq data for modified histones (H3K4me1, H3K4me2, H3K4me3, H3K9ac, H3K27ac, H3K27me3, H3K36me3, H3K79me2, and H4K20me1) were obtained from the Roadmap Epigenomics Project (39). Consolidated, mapped read data were downloaded from the Roadmap Epigenomics web portal. H3K9me3 ChIP-seq data were obtained from the ENCODE project (40). ChIP-seq data for Sororin in late S phase cells (41) and RNA-seq data from asynchronous cells (42) were downloaded from the European Nucleotide Archive (ENA) and the Sequence Read Archive (SRA), respectively. The ENCODE Repli-seq wavelet-smoothed signal was used as the replication timing score (37). The data were downloaded from the ENCODE Project portal. Details of all files used in this study are provided in Supplementary Table S4.

## Results

### ESCO2 exhibits broad chromatin binding across the genome

To characterize the genome-wide binding pattern of ESCO2, we used calibrated ChIP-seq in synchronized human cells. HeLa S3 cells were synchronized by a double-thymidine block and collected at 0 h (early S), 3 h (mid S), and 7 h (G2) after release (Fig. S1A and B). ChIP-seq was performed with an anti-ESCO2 antibody, with equal amounts of mouse chromatin spiked into each sample as an internal control to enable quantitative comparisons across conditions. Using standard peak-calling parameters, we identified fewer than 1,000 ESCO2 peaks in the human genome. This number was substantially lower than those identified for ESCO1 (∼11,800) or the cohesin subunit RAD21 (∼42,700) (Fig. S2A). Moreover, ESCO2 peaks identified in two independent mid-S experiments exhibited limited (∼15%) overlap, suggesting that many of the detected peaks are unlikely to represent genuine binding sites. These observations are consistent with previous studies reporting that ESCO2 ChIP-seq yields few, if any, clear binding peaks (20, 25).

DNA-binding proteins do not always exhibit distinct, localized peaks in ChIP-seq analyses. For example, histone H3 trimethylated at lysine 9 (H3K9me3) displays broad enrichment across large genomic regions without forming discrete peaks. We therefore examined whether ESCO2 similarly exhibits broad binding, using MUSIC, a tool that calculates the size distribution of genomic regions enriched for ChIP-seq reads (43) (Fig. S2B). As expected, the binding regions of cohesin (RAD21), acetylated cohesin (SMC3ac), and ESCO1 were typically ∼1–10 kb in width, whereas those of H3K9me3 extended over sub-megabase scales. Notably, in mid S phase, ESCO2-enriched regions also spanned hundreds of kilobases or more, supporting the view that ESCO2 displays broad chromosomal localization.

Detecting proteins with broad genomic distributions by ChIP-seq is generally challenging (except for highly abundant ones such as histones), because their occupancy probability at any given genomic site is low, leading to a poor signal-to-noise ratio. To address this, we increased the read-count bin size to 10 kb or larger, thereby averaging out stochastic variation within each bin and enabling detection even under low signal-to-noise conditions. We then used the fold enrichment (FE) between ChIP and input to correct for locus-specific variability in chromatin solubility and for known copy number variations in HeLa S3 cells. This approach allowed us to visualize sub-Mb-scale regions preferentially bound by ESCO2 during early and mid S phases (Fig. 1A). We interpret this pattern as non-localized binding, since the signal did not form discrete peaks at specific chromosomal sites. No such ESCO2-enriched domains were observed in G2 (Fig. 1A), consistent with previous findings that ESCO2 is degraded at this stage (16, 17). The ChIP-seq signal detected in G2 was likely to reflect non-specific DNA precipitation during the ChIP procedure; we therefore rescaled all FE values so that the average FE in G2 cells was set to 1 and referred to the resulting values as normalized FE (nFE). Reproducibility across two independent mid-S experiments was high, and siRNA-mediated knockdown of ESCO2 reduced nFE genome-wide to approximately 1 (Fig. S2C). Taken together, these results demonstrate that the detected sub-Mb-scale ESCO2 signals reflect genuine chromosomal association of ESCO2.

**Figure 1.**
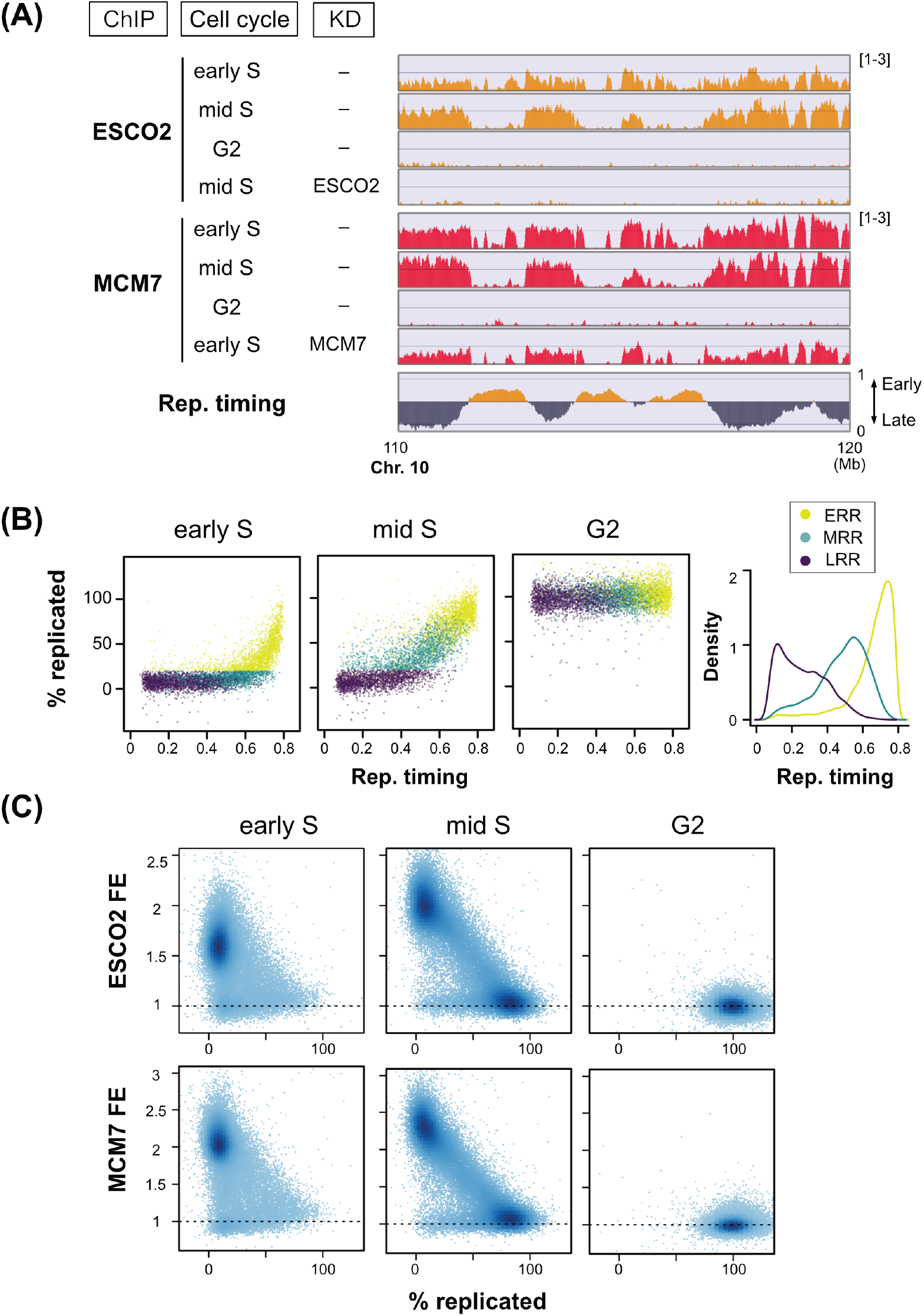
Non-localized chromosomal binding of ESCO2. (A) ChIP-seq profiles of ESCO2 and MCM7 under the indicated conditions. Broad, non-localized protein binding patterns are shown. The y-axis shows normalized fold enrichment (nFE), calculated using a 10-kb read count bin size. Replication timing (Rep. timing) is indicated by wavelet-smoothed Repli-seq signal (37). KD, knockdown by siRNA. (B) Comparison of replication timing and % replicated. Scatter plots show replication timing (x-axis) versus % replicated (y-axis), defined as the fraction of cells in which each genomic region has been replicated. Each dot represents a 100-kb region in early S, mid S, or G2. Regions initiating replication by early S, by mid S, and after mid S are defined as ERRs, MRRs, and LRRs, respectively, and are color-coded. Density plots (right) show the distributions of replication timing values for ERR, MRR, and LRR. (C) Relationship between ESCO2 or MCM7 nFE and % replicated values. Each dot represents a 100-kb genomic region.

### ESCO2 and the MCM helicase co-localize on unreplicated DNA

Human ESCO2 binding to chromosomes is mediated through its interaction with the MCM helicase (17, 20). Using calibrated ChIP-seq, we mapped the genome-wide distribution of MCM7 (a subunit of the MCM helicase) and confirmed that it similarly lacked sharp peaks, instead showing non-localized binding over sub-Mb-scale regions (Fig. 1A, S2B). A broad genomic distribution of MCM has also been reported previously in mammalian cells (44–46).

The ChIP-seq profiles showed high reproducibility between biological replicates in early S-phase cells, and knockdown of MCM7 reduced this signal, verifying that the observed distribution reflects genuine MCM7 binding (Fig. S2D). MCM7 was no longer detected on chromatin in G2 phase, consistent with the known unloading of MCM complexes from replicated chromatin (47) (Fig. 1A). A previous study reported co-localization of ESCO2 with MCM complexes in early S phase (20). Consistent with this, we observed a strong genome-wide correlation between ESCO2 and MCM7 ChIP-seq signals (r = 0.75 in early S, 0.88 in mid S) (Fig. S2E), confirming co-localization during S phase.

Different regions of the human genome replicate at characteristic time points during S phase. We found that ESCO2-and MCM7-bound regions detected in early and mid S generally corresponded to regions with late replication timing (i.e., regions with low wavelet-smoothed Repli-seq signal (37)) (Fig. 1A). To examine this relationship more closely, we computed, for each ChIP-seq sample, the fraction replicated (% replicated) at every genomic locus, which is defined as the fraction of cells in which the locus had been replicated in the populations used for ChIP-seq (Fig. 1B). Based on the resultant % replicated values in early-S, mid-S and G2 cells, we classified the entire genome into three groups: (1) early-replicating regions (ERRs), in which replication initiates by early S; (2) mid-replicating regions (MRRs), in which replication initiates by mid S; and (3) late-replicating regions (LRRs), in which replication initiates after mid S. ERRs, MRRs, and LRRs were enriched in genomic regions with high, intermediate, and low replication-timing scores derived from publicly available Repli-seq, respectively, supporting the consistency of our replication-timing classification with an independently determined replication-timing profile (Fig. 1B, right). Comparing % replicated with ESCO2 or MCM7 binding, we observed a strong inverse correlation: in fully replicated regions, the nFE signal dropped to ∼1 (background level) (Fig. 1C). These results indicate that ESCO2 and the MCM helicase specifically bind to unreplicated DNA during S phase.

### ESCO2/MCM7 binding is depleted from transcriptionally active regions

We further examined differences in ESCO2/MCM7 binding levels among unreplicated genomic regions. In early-S cells, focusing on unreplicated regions (MRR and LRR), we assessed correlations between ESCO2/MCM7 binding and various histone marks (Fig. 2A). The ChIP-seq signal showed a strong negative correlation with H3K36me3 and H3K79me2, marks associated with actively transcribed genes, as well as with RNA polymerase II (RNA pol II) binding.

**Figure 2.**
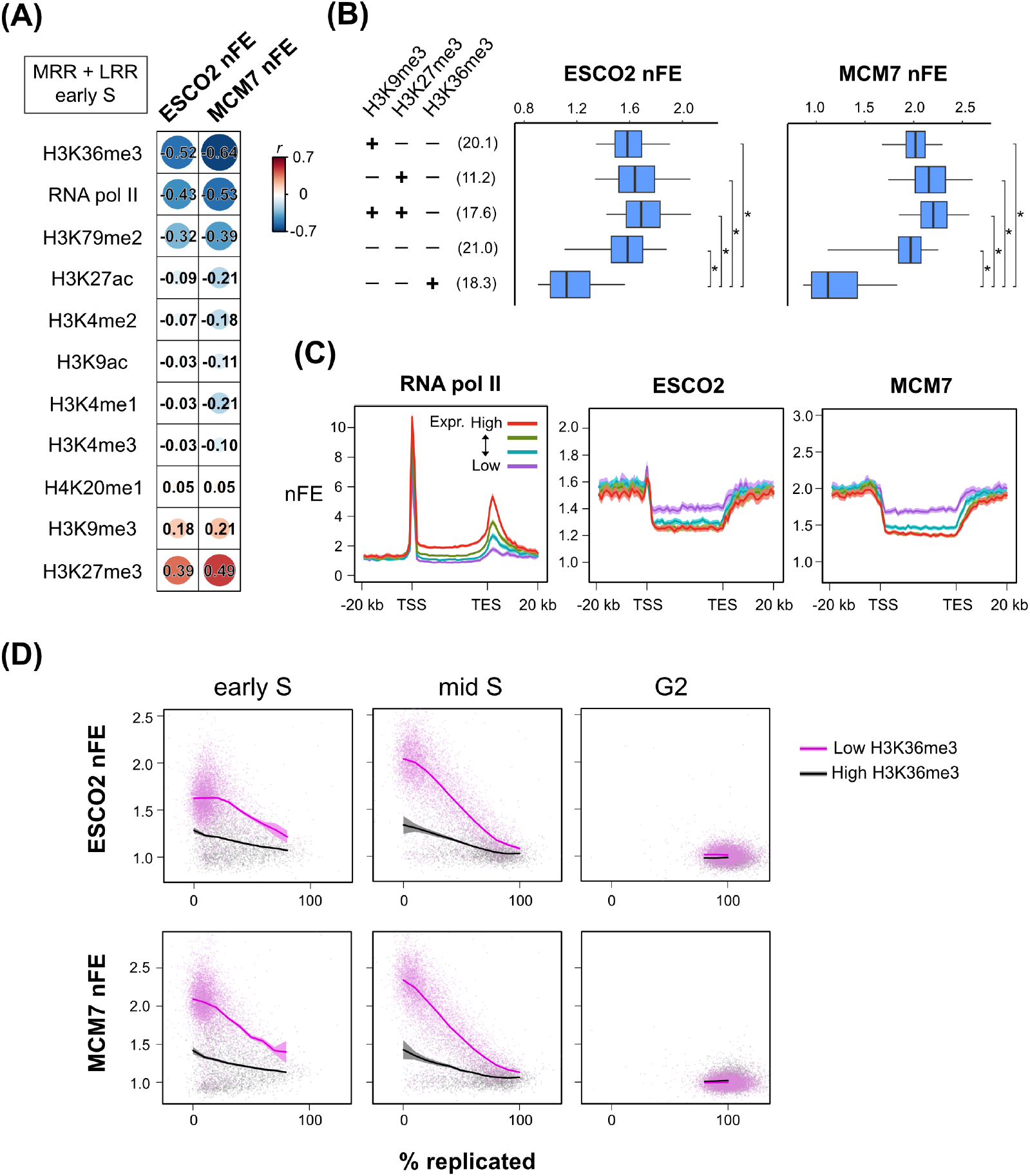
ESCO2 binds preferentially to transcriptionally inactive regions. (A) Correlation between ESCO2 or MCM7 ChIP-seq nFE and various histone modification levels or RNA polymerase II binding levels (RNA pol II). Pearson’s correlation coefficients (*r*) were calculated across the genome using 100-kb bins. Analysis was restricted to MRR and LRR in early-S cells. (B) Box plots showing the distributions of ESCO2 and MCM7 ChIP-seq nFE in early-S cells. Genomic 100-kb bins classified as MRR or LRR were subdivided into five groups according to the presence or absence of H3K36me3, H3K9me3, and H3K27me3 modifications. “+” indicates that the modification level was higher than the genome-wide average, and “−” indicates otherwise. In parentheses, the proportion (%) of bins belonging to each group relative to the whole genome is shown. Boxes indicate the interquartile range (the 25th–75th percentiles), whiskers indicate the 5th–95th percentiles, and the vertical line within each box represents the median. * *P* < 2.2 × 10⁻¹⁶, two-sided Mann–Whitney *U* test. Statistical comparisons were performed only for the indicated pairs. (C) Metagene plots of RNA pol II, ESCO2 and MCM7 ChIP-seq nFE. Genes with detectable expression were stratified into four groups according to their expression levels, and the average ChIP-seq profiles were calculated for each group. Bold lines indicate the mean, and shaded areas represent the 95% confidence interval. TSS, transcription start site; TES, transcription end site. RNA pol II data are from asynchronous cells, and ESCO2 and MCM7 data are from early-S cells. (D) Effect of H3K36me3 levels on the correlation between ESCO2 or MCM7 ChIP-seq nFE and % replicated. Bins with H3K36me3 levels below the genome-wide average are shown in magenta, and others in black. Bold lines indicate the mean, and shaded areas represent the 95% confidence interval.

Conversely, ESCO2/MCM7 binding correlated positively with heterochromatin marks such as H3K9me3 and H3K27me3.

To clarify whether ESCO2/MCM7 is depleted from transcribed regions or preferentially associated with heterochromatin, we stratified the genome into five groups based on H3K36me3, H3K27me3, and H3K9me3 levels, and compared ESCO2/MCM7 nFE across these groups (Fig. 2B). We found that nFE was low only in regions with high H3K36me3, whereas in all other groups, regardless of H3K9me3 or H3K27me3 status, it remained consistently high. This suggests depletion from transcriptionally active regions in unreplicated DNA.

Meta-gene analysis confirmed this observation. ESCO2/MCM7 binding was reduced across gene bodies, particularly in highly transcribed genes (Fig. 2C). Consistent with our findings, a recent study showed that the MCM complex is excluded from highly transcribed genes in late G1 cells (46). In Figure 2D, ESCO2/MCM7 nFE is plotted against % replicated values, with data points colored by H3K36me3 levels. Differences in ESCO2 nFE between regions with different H3K36me3 levels persisted after replication initiation and were also evident in ERR.

### ESCO1 predominantly mediates SMC3 acetylation at cohesin peaks

ESCO1 and ESCO2 both acetylate the cohesin subunit SMC3, but they are not fully redundant in establishing sister chromatid cohesion. To visualize SMC3ac localization, we synchronized HeLa S3 cells at early S, mid S, and G2 phases and performed quantitative SMC3ac ChIP-seq. We also analyzed cells subjected to siRNA knockdown of ESCO1, ESCO2, or both. Western blotting confirmed near-complete depletion of ESCO1 and/or ESCO2 (Fig. S1C). When normalized to total SMC3, SMC3ac levels measured in total cell lysates decreased moderately upon knockdown of either ESCO1 or ESCO2, while double knockdown of ESCO1 and ESCO2 caused a marked reduction (Fig. S1C, D, Fig. S3). These results are consistent with the idea that ESCO1 and ESCO2 are not fully redundant in their roles.

Calibrated ChIP-seq revealed that SMC3ac peaks in control cells co-localized with cohesin binding sites and depended on cohesin integrity, as RAD21 knockdown greatly diminished these peaks (Fig. 3A, B). Comparison of peak heights revealed little change across early S, mid S, and G2 phases (Fig. 3A, C). This observation was further validated by ChIP-qPCR analysis of SMC3ac (Fig. S4A, B). Among the knockdown conditions, ESCO2 depletion had minimal effect on peak height, whereas ESCO1 knockdown significantly reduced it, particularly in early S and mid S phases (Fig. 3C). Simultaneous depletion of ESCO1 and ESCO2 caused an even greater reduction (Fig. 3C). Consistently, ChIP-qPCR analysis of SMC3ac confirmed this trend (Fig. S4A). These findings suggest that SMC3 acetylation at cohesin peaks is primarily mediated by ESCO1, with ESCO2 providing partial compensation in the absence of ESCO1.

**Figure 3.**
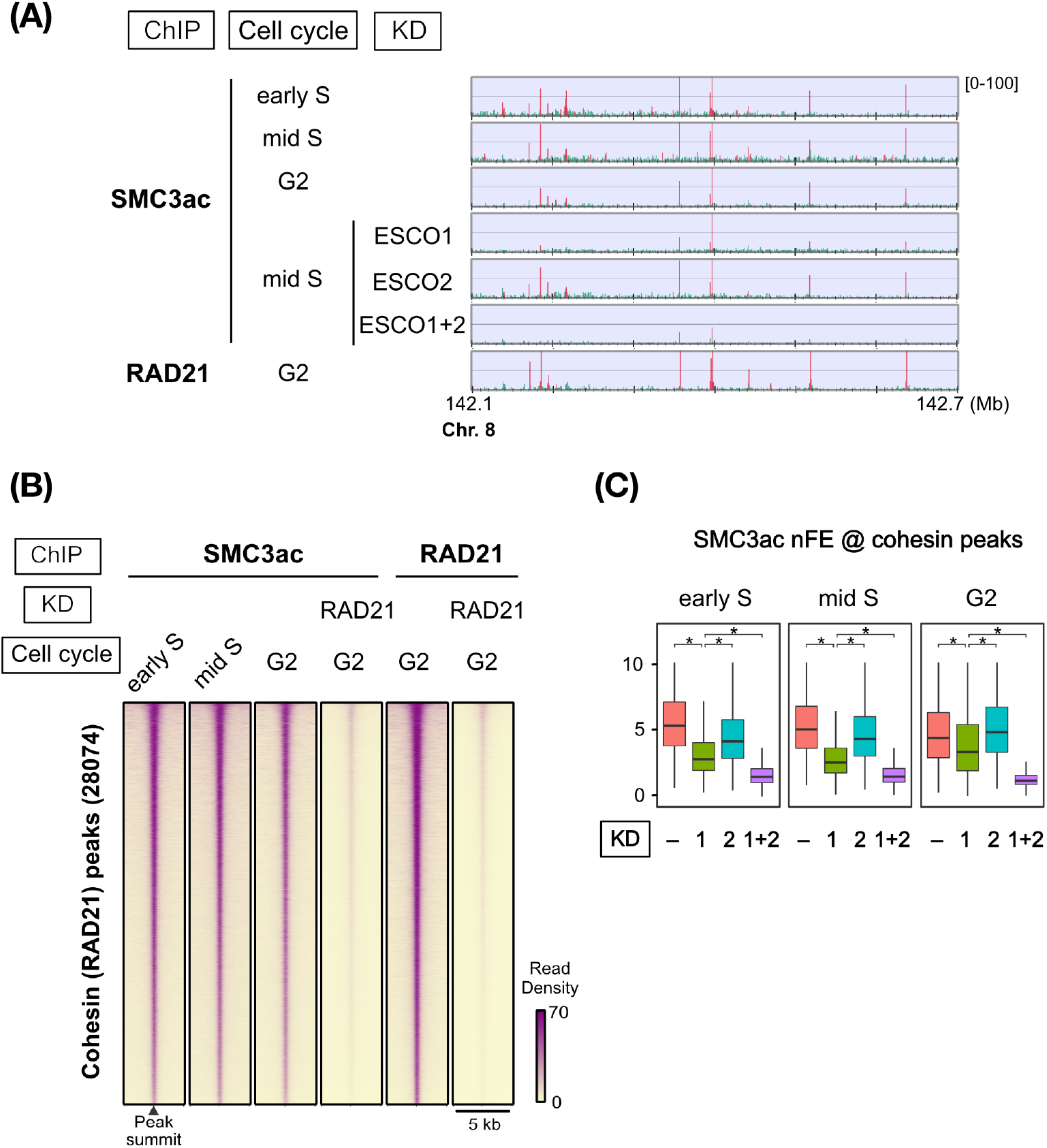
Predominant role of ESCO1 at cohesin peak sites. (A) Calibrated ChIP-seq profiles using an anti-acetyl-SMC3 antibody (SMC3ac). Read counts in the ChIP fractions at 100-bp resolution are shown, as in standard ChIP-seq analysis. Experimental conditions are indicated on the left. KD, knockdown by siRNA. ESCO1+2 indicates simultaneous KD of ESCO1 and ESCO2. RAD21 reflects cohesin localization irrespective of acetylation status. (B) Heatmaps showing SMC3ac and RAD21 ChIP-seq profiles at cohesin ChIP-seq peak sites in the genome. Experimental conditions are indicated at the top of the panel. (C) Effects of cell cycle stage and ESCO1 and/or ESCO2 knockdown on SMC3ac nFE at cohesin peak sites. KD indicates the ESCO protein(s) subjected to knockdown. Box plots show the distributions of SMC3ac nFE values under each condition. * *P* < 2.2 × 10⁻¹⁶, two-sided Mann– Whitney *U* test. Statistical comparisons were performed only for the indicated pairs.

### Differential roles of ESCO1 and ESCO2 in non-localized cohesin acetylation

Given that cohesin at ChIP-seq peaks is not the primary target of ESCO2-dependent acetylation, we investigated whether ESCO2 acetylates cohesin bound to chromatin outside peak regions. To this end, we divided the genome into 100 bp bins, excluded bins corresponding to cohesin peak regions, and computed nFE values by aggregating the remaining bins at 100-kb resolution. Hereafter, “SMC3ac nFE” refers to this non-localized signal, representing SMC3 acetylation outside cohesin peaks. In Figure 4A, SMC3ac nFE is plotted against % replicated values. In control HeLa cells in early and mid S, nFE increased with replication progress, consistent with replication-coupled acetylation of non-localized cohesin. Notably, SMC3ac nFE peaked when % replicated reached ∼50% and then declined, suggesting that newly generated SMC3ac is not stably retained on chromosomes over time (see below for details).

**Figure 4.**
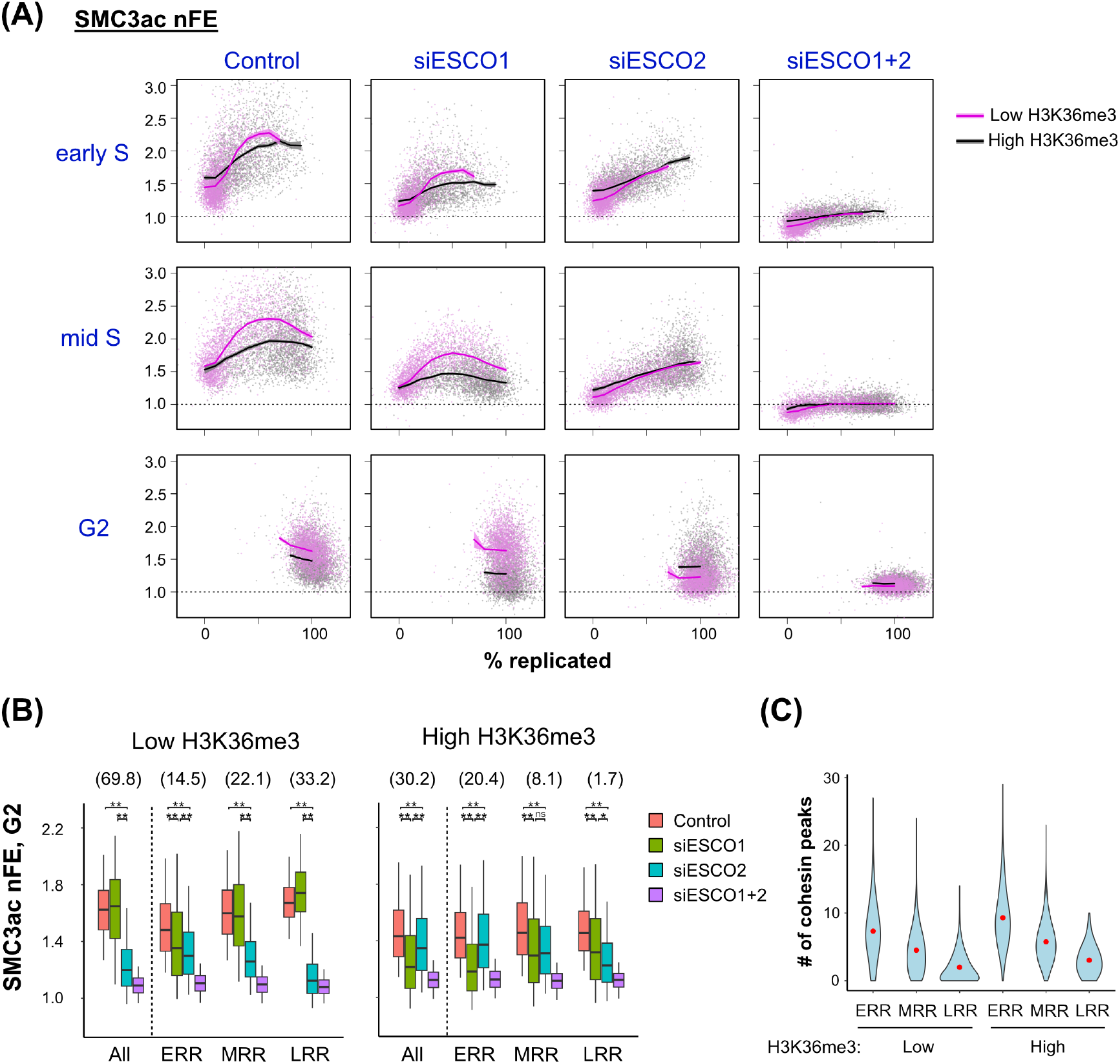
Differential roles of ESCO1 and ESCO2 in non-localized cohesin acetylation. (A) Scatter plots showing the relationship between non-localized SMC3ac ChIP-seq nFE and % replicated. Cells were treated with siESCO1, siESCO2, or both (siESCO1+2), synchronized at early S, mid S, or G2, and then analyzed. Control indicates mock transfection. Each dot represents a 100-kb genomic bin. Bins with H3K36me3 levels below the genome-wide average are shown in magenta, and others in black. Bold lines indicate the mean, and shaded areas represent the 95% confidence interval. (B) Box plots showing non-localized SMC3ac ChIP-seq nFE in G2-phase cells. Genomic 100-kb bins were classified into six groups according to H3K36me3 levels and replication timing, and the distributions of nFE in each group are shown. “Low” and “high” H3K36me3 indicate genomic regions with H3K36me3 levels below and above the genome-wide average, respectively. “All” indicates the combined set of ERR, MRR, and LRR. The proportion (%) of bins belonging to each group is shown in parentheses. Knockdown conditions are distinguished by color. Boxes represent the interquartile range (the 25th–75th percentiles), whiskers indicate the 5th–95th percentiles, and the horizontal line within each box denotes the median. Statistical comparisons were performed for the indicated pairs (two-sided Mann–Whitney *U* test). **, *P* < 2.2 × 10⁻¹⁶; *, *P* = 1.5 × 10⁻^5^; ns, *P* > 0.01. (C) Violin plots showing the distribution of cohesin peak numbers. As in (B), genomic 100-kb bins were classified into six groups. For each bin, the number of cohesin (RAD21) ChIP-seq peaks within a 500-kb window centered on the midpoint of the bin was calculated. Red dots indicate the mean.

We next examined the dependence of replication-coupled SMC3 acetylation on ESCO1 and ESCO2. In cells with combined ESCO1 and ESCO2 knockdown, the replication-associated increase in SMC3ac nFE was almost completely abolished, whereas depletion of either enzyme alone resulted in an attenuated increase (Fig. 4A). These results indicate that both ESCO1 and ESCO2 contribute to the acetylation of non-localized cohesin. However, the relative contribution of ESCO1 and ESCO2 varied across genomic regions. Regions with low H3K36me3, which account for approximately 70% of the genome, showed a greater increase in SMC3ac levels during S phase when ESCO2 was present (control or ESCO1 knockdown cells), consistent with the preferential localization of ESCO2 to these regions (Fig. 4A). Even in G2-phase cells, in which overall SMC3ac nFE had declined, low-H3K36me3 regions retained relatively higher SMC3ac levels in an ESCO2-dependent manner (Fig. 4A, B).

In contrast, we found that the contribution of ESCO1 varied markedly depending on the replication timing of genomic domains. The contribution of ESCO1 to non-localized SMC3ac levels in G2 was greater in early-replicating regions (ERR) than in MRR or LRR, regardless of histone modification status (Fig. 4B). This tendency was particularly pronounced within ERR regions enriched for H3K36me3, where ESCO2 binding is scarce and ESCO1 emerged as the primary contributor to non-localized SMC3ac generation. By contrast, in MRR and LRR with low H3K36me3 levels, which account for approximately 55% of the genome, the reduction in SMC3ac upon ESCO2 knockdown was nearly identical to that observed in cells with combined ESCO1 and ESCO2 knockdown (Fig. 4B), indicating that ESCO2 serves as the major enzyme responsible for non-localized SMC3 acetylation in these regions. ESCO1 has been shown to co-localize with cohesin peaks (18). We also found that the density of cohesin peaks was higher in ERR than in MRR or LRR (Fig. 4C). Given that cohesin rings can passively diffuse along DNA (48), these results suggest a model in which ESCO1-dependent non-localized SMC3ac, enriched in ERR, arises from ESCO1-mediated acetylation at cohesin peak sites followed by diffusion of acetylated cohesin along the DNA.

### HDAC8-dependent decline of non-localized SMC3ac after its generation in S phase

SMC3ac levels do not continuously increase throughout S phase, leading us to track how SMC3ac levels change after the completion of DNA replication. We collected late G2 cells by culturing cells released from a thymidine block for 8.5 h, which is 1.5 h longer than the earlier G2 time point, and performed quantitative SMC3ac ChIP-seq. When comparing late G2 with early S, mid S, and G2, we observed that in control cells, SMC3ac levels peaked in early and mid S for ERR, in mid S for MRR, and in G2 for LRR, consistent with their replication timing (Fig. 5A). Importantly, SMC3ac levels ceased to increase across all domain classes (ERR, MRR, and LRR) once replication was completed and instead declined thereafter. Thus, replication-coupled SMC3ac is not stably maintained on chromatin once replication is completed. These observations indicate that the dynamics of non-localized SMC3ac are governed by two opposing processes, namely its generation during replication and its subsequent loss, both of which must be considered to explain its overall behavior.

**Figure 5.**
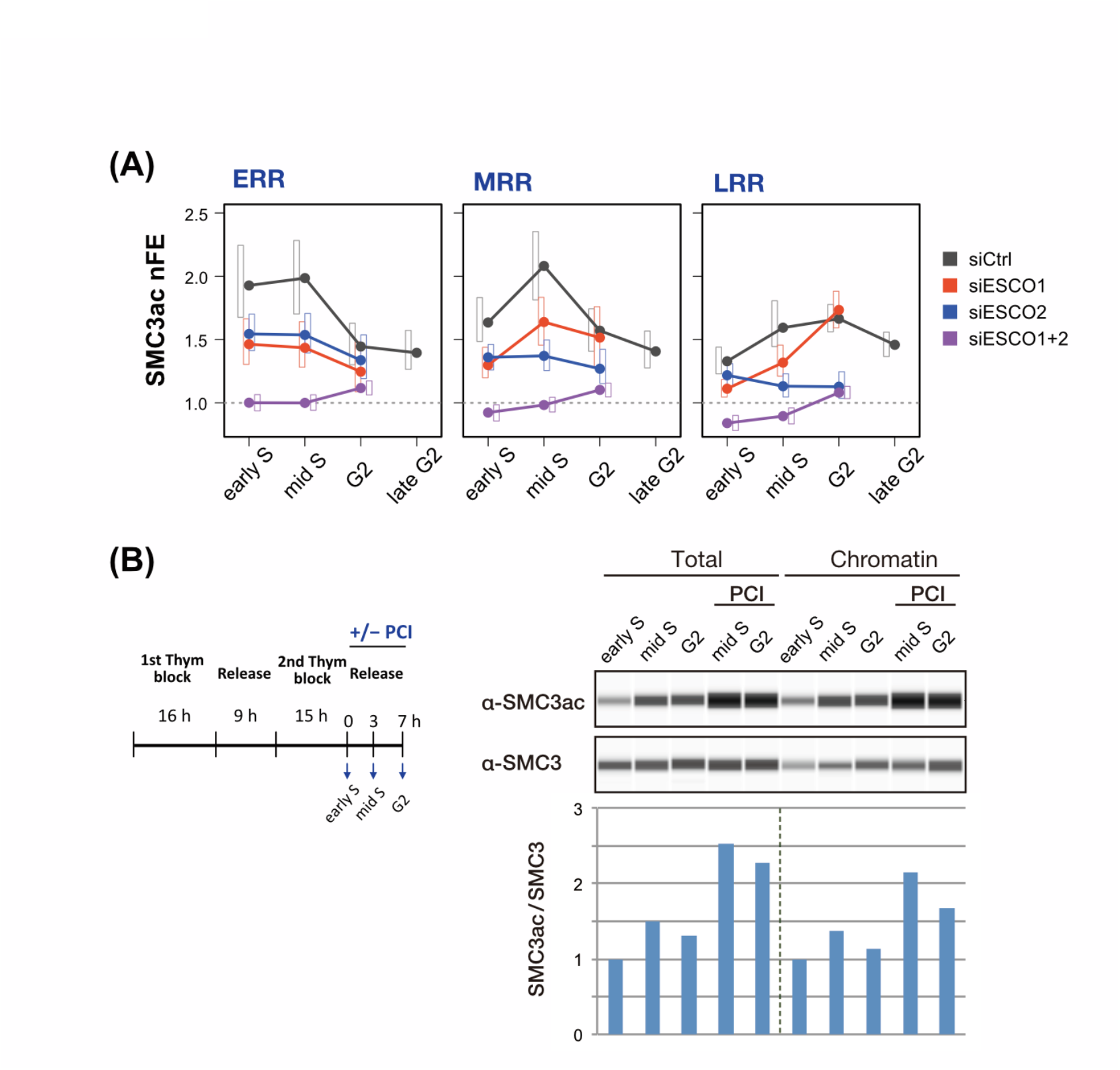
HDAC8-dependent decline of non-localized SMC3ac after its generation in S phase. (A) Dynamics of non-localized SMC3ac ChIP-seq nFE across the cell cycle in ERR, MRR, and LRR. Knockdown conditions are distinguished by color. Dots indicate the median, and boxes represent the interquartile range. For the control (mock-treated) sample, data from cells cultured 1.5 h longer than G2-phase samples (late G2) are also shown. (B) Effects of the HDAC8 inhibitor PCI-34051 on chromosomal SMC3ac levels. (Left) Cell culture conditions. Thym, thymidine; PCI, PCI-34051. (Right) Quantification of SMC3ac and SMC3 levels. Total extracts and chromatin fractions prepared under the indicated conditions were analyzed using an automated capillary western system and presented as virtual blot-like images. Ratios of SMC3ac to SMC3 are shown below as bar graphs. Values were rescaled so that those in early S were set to 1.

Histone deacetylase 8 (HDAC8) is known to catalyze SMC3 deacetylation (28, 49). We tested whether HDAC8 contributes to the observed loss of SMC3ac. HeLa S3 cells synchronized by thymidine release were split into two groups, one of which was treated with the HDAC8 inhibitor PCI-34051 during S-phase progression (Fig. 5B, left). Cells were collected at 0, 3, and 7 h (corresponding to early S, mid S, and G2). Total lysate and chromatin fractions were prepared and analyzed by quantitative western blotting with anti-SMC3 and anti-SMC3ac antibodies. The SMC3ac/SMC3 ratio was used as a measure of acetylation level. In untreated cells, acetylation increased from early to mid S and then declined by G2, consistent with the trends for non-localized cohesin acetylation observed by calibrated ChIP-seq (Fig. 5B, right). In inhibitor-treated cells, acetylation levels in mid S and G2 were significantly higher than in untreated cells, and this effect was observed in both total lysate and chromatin fractions (Fig. 5B, right). These results independently support the reduction of chromatin-bound SMC3ac observed in ChIP-seq and indicate that this decrease is at least partly dependent on HDAC8.

### SMC3ac is preferentially retained in H3K27me3-rich chromatin

We next asked whether the loss of SMC3ac occurs uniformly across the genome. For various histone modifications and RNA pol II binding, we extracted genomic regions with low (bottom 25%) or high (top 25%) signal levels and quantified differences in SMC3ac levels between them using Cohen’s d (Fig. 6A). In G2 cells (shortly after replication completion), SMC3ac showed negative associations with H3K36me3 and RNA polymerase II binding, as well as with several other histone marks enriched in transcribed regions. These associations likely reflect both reduced ESCO2 binding in transcriptionally active regions and the enrichment of such regions in ERRs (Fig. S5), where SMC3ac loss has progressed further by G2. In contrast, H3K27me3 exhibited a strong positive association with SMC3ac in G2 cells (Fig. 6A).

**Figure 6.**
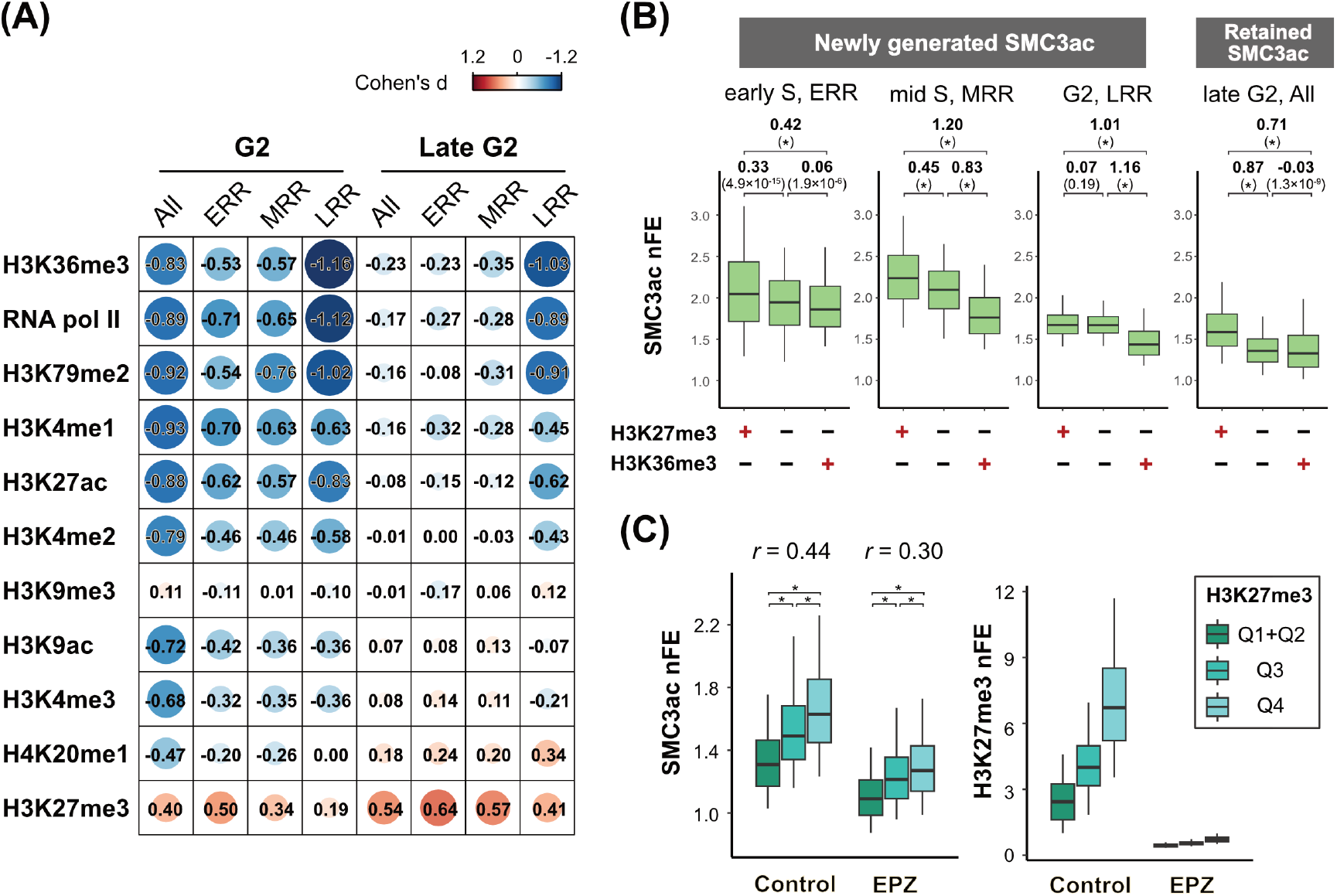
Preferential retention of SMC3ac in H3K27me3-marked regions. (A) Relationship between histone modifications and SMC3ac binding levels. The genome was divided into 100-kb bins, and differences in SMC3ac binding between the top and bottom quartiles (each 25%) of each histone modification level were evaluated using Cohen’s d. RNA pol II indicates RNA polymerase II binding. SMC3ac levels were measured in mock-treated cells at G2 and late G2, as indicated. Data are presented for the whole genome (All) as well as for ERR, MRR, and LRR regions classified by replication timing. (B) SMC3ac nFE in genomic regions with different H3K27me3 and H3K36me3 levels. The genome was divided into 100-kb bins, which were classified into three categories: bins with only H3K27me3 levels above the genome-wide average, bins with only H3K36me3 levels above the genome-wide average, and bins with both H3K27me3 and H3K36me3 levels at or below their respective genome-wide averages. These three categories together accounted for 95.5% of the genome (Fig. S6A). The distribution of SMC3ac nFE in each category is shown as boxplots. Boxes indicate the interquartile range (25th–75th percentiles), whiskers indicate the 5th to 95^th^ (Fig. 6 legend, continued) percentiles, and thick horizontal lines indicate the medians. The left three panels show SMC3ac distributions shortly after its generation, focusing on ERRs in early S phase, MRRs in mid S phase, and LRRs in G2, respectively. The rightmost panel shows retained SMC3ac levels across all genomic regions in late G2. Effect sizes for pairwise comparisons were assessed using Cohen’s *d* and are shown in bold. *P* values from two-sided Mann–Whitney *U* tests are shown in parentheses. *, *P* < 2.2 × 10⁻¹⁶. (C) Association between chromosomal SMC3ac levels and H3K27me3 and the effect of EZH2 inhibition. Based on H3K27me3 levels from ENCODE data for untreated cells, the genome was divided into three groups (Q1+Q2: below the median, Q3: between the median and the third quartile, Q4: above the third quartile). Distributions of SMC3ac ChIP-seq nFE in late G2 cells, with or without EZH2 inhibitor treatment, were compared across these groups. H3K27me3 levels measured under the same conditions are also shown to assess the effect of EZH2 inhibition. Boxes represent the interquartile range (25th–75th percentiles), whiskers indicate the 5th–95th percentiles, and bold lines denote the median. Data are shown for the whole genome (All) as well as for ERR, MRR, and LRR regions classified by replication timing. Pearson correlation coefficients between SMC3ac and H3K27me3 levels are indicated as *r*. * *P* < 2.2 × 10⁻¹⁶, two-sided Mann–Whitney *U* test.

In late G2 (a later time point after replication completion), the association with H3K27me3 became even stronger, whereas associations with H3K36me3 and other transcription-associated marks largely disappeared (Fig. 6A). Stratification by replication timing revealed that the positive association between SMC3ac and H3K27me3 was more pronounced in ERR and MRR than in LRR (Fig. 6A, B). Conversely, correlations with histone marks other than H3K27me3 largely disappeared in ERR and MRR but persisted in LRRs (Fig. 6A), consistent with the shorter time elapsed since SMC3ac generation in LRRs and partial preservation of its initial distribution. Genomic regions could be broadly classified into three mutually exclusive groups: H3K27me3-rich regions, H3K36me3-rich regions, and regions with low levels of both modifications (Fig. S6A). In late G2, SMC3ac nFE was higher in H3K27me3-rich regions than in either of the other two groups (Fig. 6B, right panel). In contrast, shortly after SMC3ac generation, particularly in MRRs and LRRs where ESCO2 plays a predominant role in SMC3 acetylation, the major feature was notably reduced SMC3ac nFE in H3K36me3-rich regions (Fig. 6B, two middle panels). Thus, the enrichment of SMC3ac in H3K27me3-rich regions observed in late G2 cannot simply be explained by its distribution at the time of SMC3ac generation and instead suggests preferential retention of SMC3ac in H3K27me3-rich chromatin.

To validate these observations, we first examined the reproducibility of the late-G2 SMC3ac ChIP-seq profiles between biological replicates. The replicates showed substantial genome-wide agreement (Fig. S6B), and the positive correlation between SMC3ac nFE and H3K27me3 levels was reproduced in the independent replicate (Fig. 6C, left panel, control). We next performed SMC3ac ChIP-qPCR at representative loci. Genomic loci enriched for H3K27me3 consistently exhibited higher SMC3ac enrichment than loci with low H3K27me3 levels, in agreement with the ChIP-seq results (Fig. S6C). SMC3ac signals were reduced at all loci following combined depletion of ESCO1 and ESCO2. Together, these results indicate that acetylated cohesin is broadly distributed across the genome but preferentially enriched at H3K27me3-rich regions.

Sororin preferentially associates with acetylated cohesin (14, 50). We therefore asked whether Sororin exhibits similar chromatin dynamics by conducting calibrated Sororin ChIP-seq from early S through late G2. Sororin was enriched at cohesin peak regions (Fig. S7A), consistent with previous observations (41). Its peak-associated signal increased during S phase (Fig. S7A, B), consistent with the accumulation of Sororin protein during S phase (50). Non-localized Sororin signals similarly increased during S phase, and both peak-associated and non-localized Sororin signals declined markedly by late G2 (Fig. S7B), broadly resembling the dynamics of SMC3ac. Non-localized Sororin also showed a positive association with H3K27me3 in G2 (Fig. S7C, left), and a similar association was observed in an independent, previously published Sororin ChIP-seq dataset from late S-phase cells (Fig. S7C, right). In late G2, however, the Sororin ChIP-seq signal-to-noise ratio was substantially lower (Fig. S7A), and a clear association with H3K27me3 was not observed. Consistent with a substantial contribution of background signal under this condition, ESCO2 depletion did not reduce the apparent non-localized Sororin signal in late G2 (Fig. S7B). Thus, although Sororin dynamics broadly mirrored those of SMC3ac, its association with H3K27me3 in late G2 could not be reliably assessed with the current dataset.

Finally, we examined whether H3K27me3 contributes to this enrichment. We treated cells with the selective EZH2 inhibitor EPZ-6438 (51), which almost completely abolished genome-wide H3K27me3 signals (Fig. 6C). Under these conditions, non-localized SMC3ac signals were also reduced, with the greatest decreases occurring in genomic regions that exhibited the highest H3K27me3 levels in untreated cells (Fig. 6C). Accordingly, the correlation between non-localized SMC3ac and H3K27me3 decreased from 0.44 to 0.30 following inhibitor treatment (Fig. 6C). These observations support the idea that H3K27me3, or the chromatin environment associated with H3K27me3, contributes to the preferential retention of non-localized SMC3ac in late G2. However, the reduction in SMC3ac was substantially smaller than the nearly complete loss of H3K27me3 itself.

## Discussion

In this study, we uncovered two previously unrecognized layers in the generation of ESCO2-mediated cohesion: the broad, replication-coupled acetylation of non-peak cohesin across the genome, and the region-specific retention of SMC3ac within H3K27me3-rich domains. These findings refine current models of how cohesin acetylation is generated and stabilized, and they shed new light on the genome-wide architecture of sister chromatid cohesion.

Our quantitative ChIP-seq analyses revealed that ESCO2 binds preferentially to unreplicated, transcriptionally inactive regions during S phase (Figs. 1, 2). Unlike sharp, discrete peaks generated by factors such as CTCF, ESCO2 exhibited a broad distribution extending over hundreds of kilobases. This binding mode likely explains why previous efforts to map ESCO2 genomic occupancy consistently failed to detect clear peaks (20, 25). Consistent with its binding pattern, the SMC3 acetylation catalyzed by ESCO2 also showed a diffuse genomic distribution (Fig. 4). Although biochemical studies have established that cohesin acetylation increases during S phase (28, 41), it remained unclear where in the genome these modifications occur, because SMC3ac levels at canonical cohesin peaks remained unchanged throughout S phase (Fig. 3). Our data reconcile this discrepancy by demonstrating that the majority of replication-coupled acetylation occurs on non-peak cohesin distributed broadly along chromosomes.

Because both ESCO2 and SMC3ac signals were only marginally above background, calibrated ChIP-seq required careful minimization of stochastic variation. Increasing the bin size to 100 kb greatly improved the signal-to-noise ratio while maintaining meaningful spatial resolution, as this scale remains smaller than the size of a typical TAD (52). This analytical approach may be useful for profiling other chromatin-associated factors with similarly non-localized binding patterns.

ESCO2 is known to interact with the replicative helicase MCM (17, 20), and their ChIP-seq profiles were nearly identical (Figs. 1, S2). Importantly, ESCO2 was detected in mid-and late-replicating regions even before replication had initiated in these domains (Fig. 1), indicating that ESCO2 associates with pre-replication complexes prior to origin firing. This observation is consistent with data from *Xenopus* egg extracts (19). Furthermore, our data show that cohesin acetylation within a given genomic region progresses in parallel with replication of that region (Fig. 5A). A large number of MCM complexes are loaded onto chromatin during G1, whereas only a small fraction of them are eventually activated as replicative helicases (53). In addition, *in vitro* reconstitution studies have shown that activated replicative helicases can push unfired MCM complexes ahead of the progressing fork (54). In this study, we found that MCM-ESCO2 complexes accumulate preferentially in transcriptionally inactive regions (Fig. 2). Taken together, these observations suggest that, in such regions, ESCO2 may be recruited in greater amounts through its association with unfired, fork-proximal MCM complexes. Consequently, cohesin encountered by the replication machinery in transcriptionally inactive regions may undergo SMC3 acetylation more efficiently.

A major discovery of this study is that SMC3 acetylation generated during S phase is not stably maintained but gradually diminishes over time (Fig. 5). The fraction that persists into late G2 is strikingly enriched in genomic regions with high levels of H3K27me3 (Fig. 6). Because the presence of cohesive cohesin can interfere with promoter-enhancer communication and transcriptional elongation, its selective stabilization in transcriptionally repressed domains may represent an evolved strategy to limit the disruptive potential of cohesive cohesin at active genes.

The preferential retention of SMC3ac in H3K27me3-rich chromatin does not appear to be simply a consequence of low transcriptional activity. In late G2, SMC3ac levels were higher in H3K27me3-rich regions than in regions with low levels of both H3K27me3 and the transcription-associated mark H3K36me3 (Fig. 6B). Moreover, depletion of H3K27me3 by EZH2 inhibition reduced non-localized SMC3ac, with the greatest reductions occurring in regions that had the highest H3K27me3 levels before inhibitor treatment (Fig. 6C). These observations suggest that H3K27me3, or a chromatin environment associated with H3K27me3, contributes to the preferential retention of acetylated cohesin. However, the reduction in SMC3ac was considerably smaller than the nearly complete loss of H3K27me3 following EZH2 inhibition. Thus, the preferential retention of SMC3ac is unlikely to depend simply on direct recognition of H3K27me3 by acetylated cohesin. Determining the molecular basis of this preferential retention will be an important subject for future studies.

Roberts syndrome, caused by ESCO2 dysfunction, is characterized by developmental defects, growth retardation, and chromosome segregation abnormalities (55, 56). Cells from affected individuals display cohesion loss near centromeres. Mammalian pericentromeric regions have been reported to contain H3K27me3-marked chromatin (57, 58), raising the possibility that the preferential maintenance of SMC3ac in such chromatin contributes to robust pericentromeric cohesion. However, pericentromeric regions are highly repetitive and are therefore poorly represented in conventional short-read sequencing data such as those used in this study. Consequently, our results do not exclude the possibility that ESCO2-generated SMC3ac is also stably maintained in H3K9me3-rich pericentromeric chromatin through mechanisms distinct from those associated with H3K27me3-rich regions. Defective cohesion between sister kinetochores is believed to impair faithful chromosome segregation. Indeed, patient-derived cells exhibit mitotic delay, aberrant mitotic figures, and reduced proliferative capacity, which are thought to underlie developmental defects (59). The current study highlights the importance of chromatin context in shaping the functional outcomes of ESCO2-dependent cohesin acetylation and provides a mechanistic framework for understanding cohesion defects in human disease.

## Acknowledgements

We thank all members of the Shirahige laboratory for helpful discussions and continuous support. We also thank Prof. Yukiko Okada for providing cells from the bone marrow and spleen of H2B-GFP mice.

## Data availability

All ChIP-seq data generated in this study have been deposited in the Gene Expression Omnibus (GEO) under the accession number GSE312990.

## Author contributions

Conceptualization: K.S., T.Su.

Methodology: M.I.S., T.Su.

Investigation: M.I.S., M.B., A.Y., T.Sa.

Resources: T.N.

Data curation: M.I.S.

Formal analysis: M.I.S., T.Su.

Visualization: M.I.S., T.Su.

Writing – original draft: T.Su.

Writing – review & editing: All authors

Supervision: K.S., T.Su.

Project administration: K.S., T.Su.

Funding acquisition: M.I.S., K.S., T.Su.

## Use of generative AI

Generative artificial intelligence was used for language editing of the manuscript; all scientific content and interpretations were produced by the authors.

