## Supplemental Materials for "Region-specific generation and retention of acetylated cohesin shape the genome-wide pattern of sister chromatid cohesion"

### Figure S1

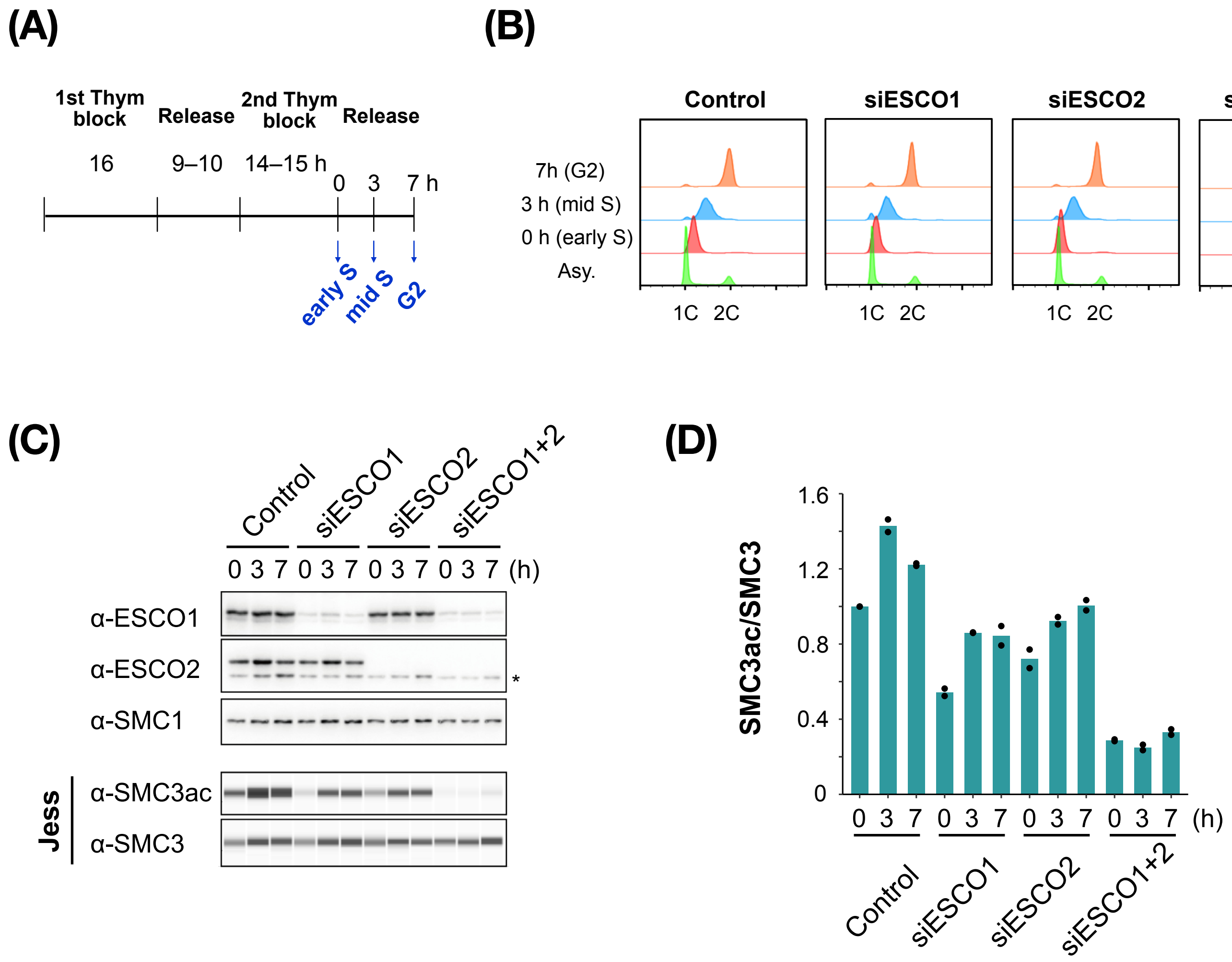


**Figure S1 Cell culture validation**

(A) Scheme of cell synchronization.

(B) Verification of synchronization by flow cytometry. Cells were transfected with siRNA targeting ESCO1 (siESCO1), ESCO2 (siESCO2), or both (siESCO1+2). Representative results from multiple independent experiments are shown. Asy, asynchronous.

(C) Validation of siRNA knockdown of ESCO1 and ESCO2. Protein levels of ESCO1 and ESCO2 were examined by western blotting, with the cohesin subunit SMC1 as a loading control. SMC3 acetylation levels were measured using an automated capillary western system (JESS) and presented as virtual blot images. Total SMC3 levels were also measured. Representative results from repeated experiments are shown.

(D) Ratios of SMC3ac to SMC3 measured by JESS. Results from two independent experiments are shown as dots, with the mean shown as a bar.

### Figure S2

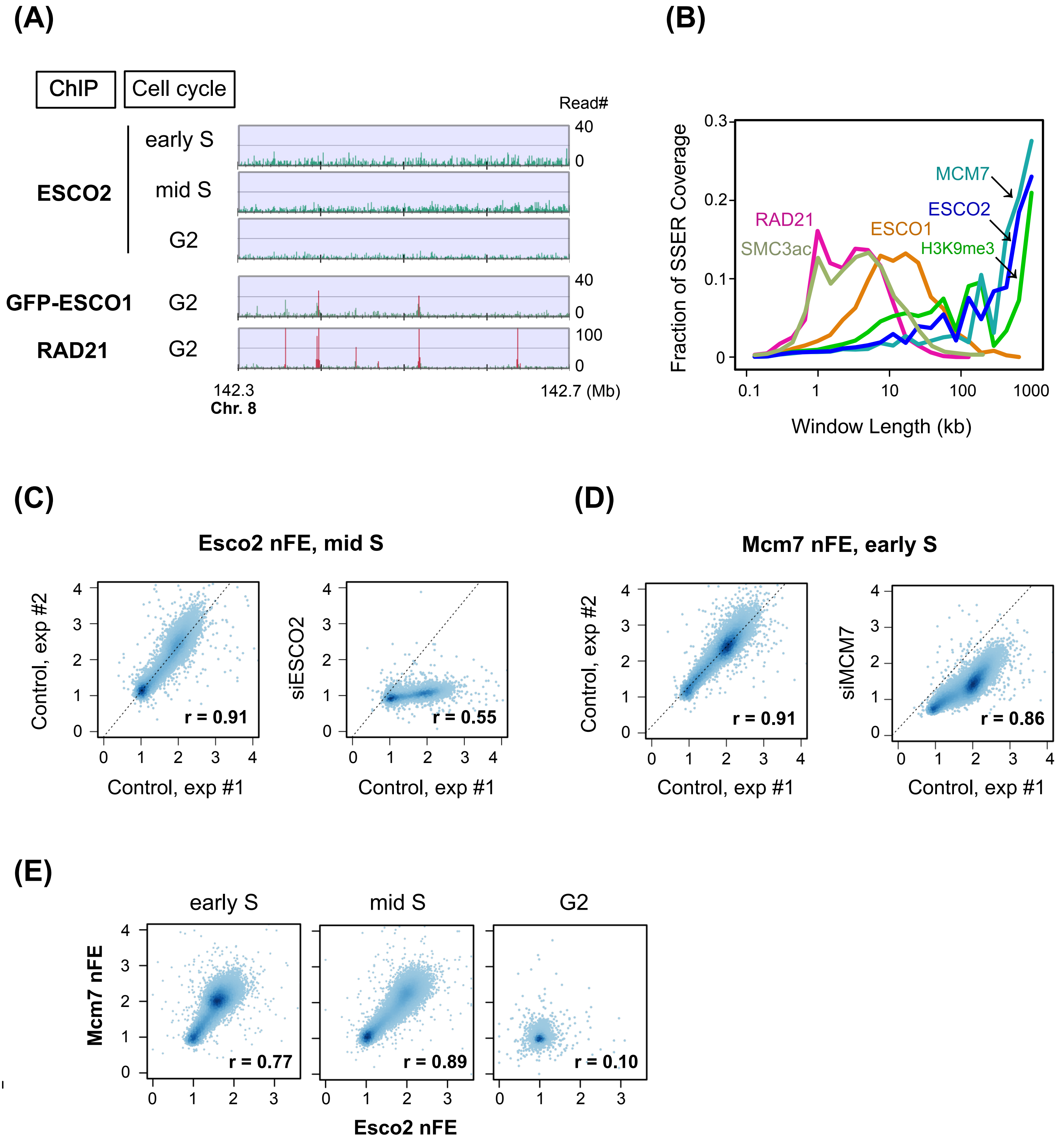

(A)
ChIP
Cell cycle
Read#
ESCO2
early S
mid S
G2
GFP-ESCO1
G2
RAD21
G2
40
0
40
0
100
0
142.3
Chr. 8
142.7 (Mb)
(B)
Fraction of SSER Coverage
Window Length (kb)
MCM7
ESCO2
RAD21
ESCO1
SMC3ac
H3K9me3
(C)
Esco2 nFE, mid S
Control, exp #2
siESCO2
Control, exp #1
r = 0.91
r = 0.55
(D)
Mcm7 nFE, early S
Control, exp #2
siMCM7
Control, exp #1
r = 0.91
r = 0.86
(E)
early S
mid S
G2
Mcm7 nFE
Esco2 nFE
r = 0.77
r = 0.89
r = 0.10

**Figure S2 Supplementary data for ESCO2 and MCM7 ChIP-seq analysis**

(A) ChIP-seq profiles of ESCO2, GFP-ESCO1, and RAD21 at 100-bp resolution. Experimental conditions are indicated on the left. Detected peaks are highlighted in red.

(B) Distribution of enriched region sizes in ChIP-seq data for the indicated proteins, analyzed using MUSIC. SSER, scale-specific enriched region. RAD21 and SMC3ac data are from G2 cells, ESCO2 data are from mid-S cells, and MCM7 data are from early-S cells.

(C) Validation of ESCO2 ChIP-seq data. Scatter plots show high correlations of ChIP-seq nFE values between two biological replicates of untreated controls (left) and a reduction of nFE upon ESCO2 knockdown (right). Each dot represents a 100-kb bin. Data are from mid-S cells. $r$, Pearson's correlation coefficient.

(D) Validation of MCM7 ChIP-seq data. Scatter plots show high correlations of ChIP-seq nFE values between two biological replicates of untreated controls (left) and a reduction of nFE upon MCM7 knockdown (right). Each dot represents a 100-kb bin. Data are from early-S cells. $r$, Pearson's correlation coefficient.

(E) Comparison of ESCO2 and MCM7 ChIP-seq profiles across early S, mid S, and G2. ChIP-seq nFE was calculated at 100-kb resolution. $r$, Pearson's correlation coefficient.

### Figure S3

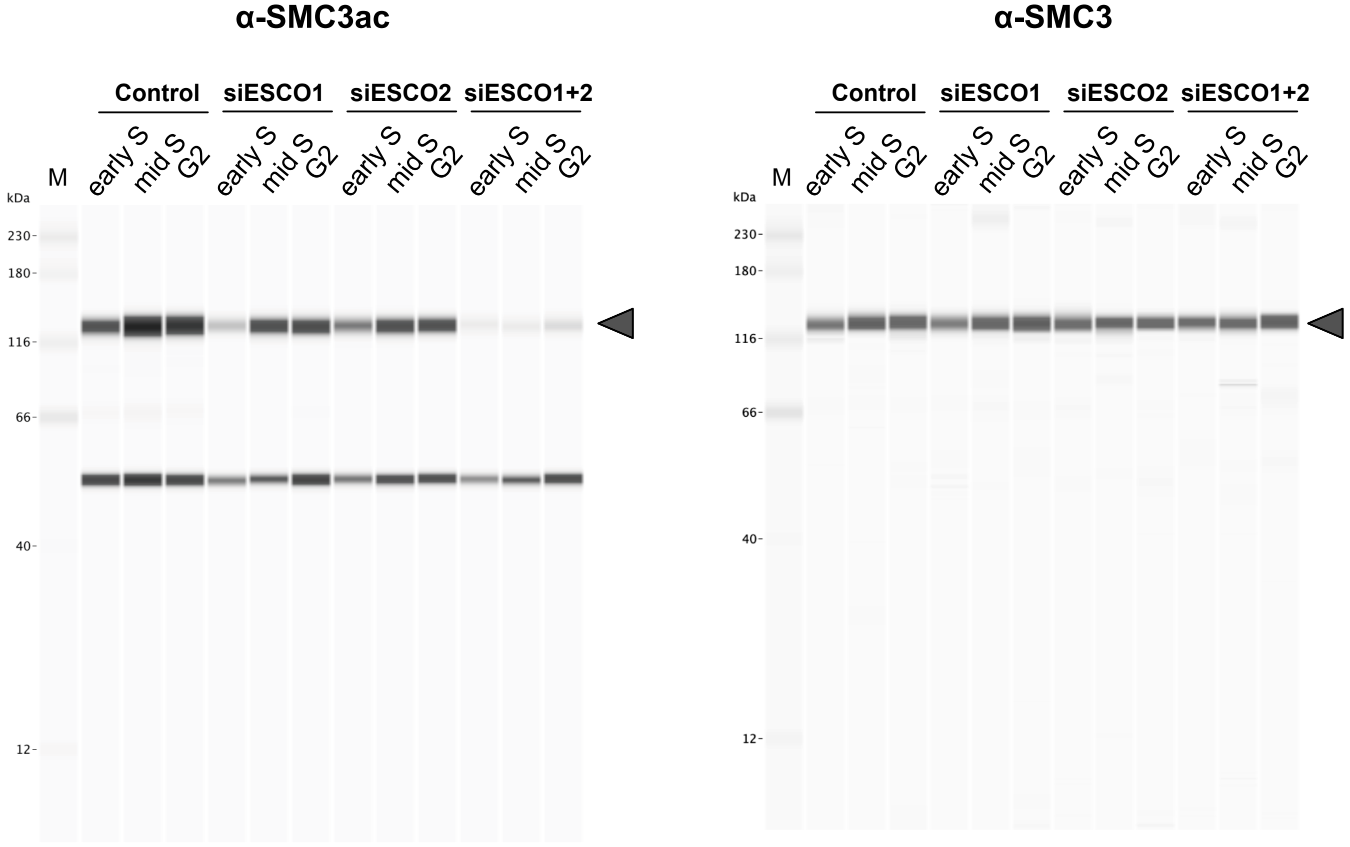


**Figure S3 Antibody validation for automated capillary western system**
Uncropped data corresponding to Fig. S1C. The anti-SMC3ac and anti-SMC3 antibodies detected signals at the same molecular weight (gray arrowheads), consistent with the expected size of SMC3 (~142 kDa).

### Figure S4

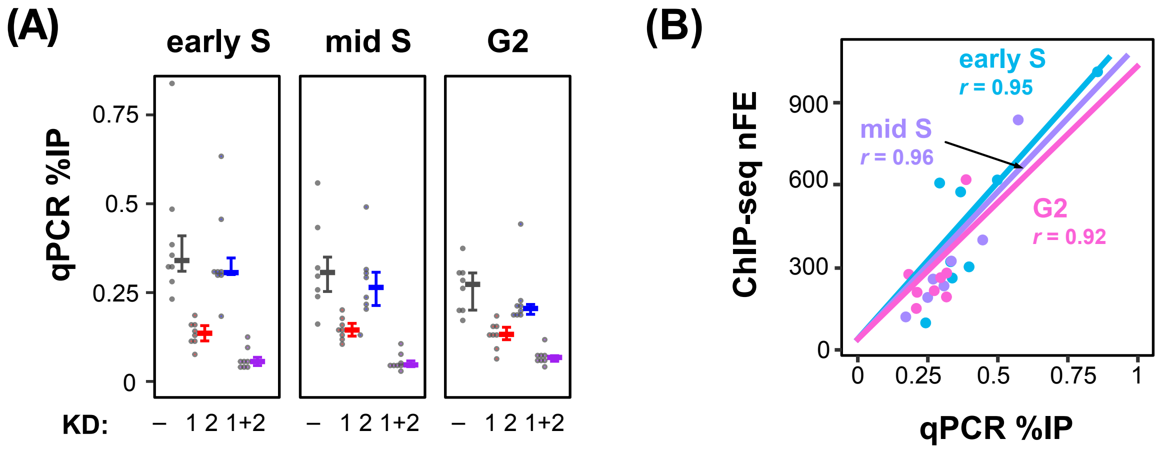


**Figure S4 Validation of SMC3ac ChIP-seq data by qPCR**

(A) ChIP efficiency of SMC3ac at eight cohesin peak sites quantified by qPCR. KD indicates the targeted ESCO protein(s) by siRNA. Error bars represent the interquartile range, and horizontal bold lines indicate the median.

(B) Comparison of SMC3ac ChIP efficiency measured by qPCR and ChIP-seq peak heights at the same eight cohesin peak sites in control cells without knockdown. Data from early S, mid S, and G2 cells each showed strong correlations, with similar slopes of the regression lines. These results demonstrate appropriate calibration of ChIP-seq data using spike-in controls.

### Figure S5

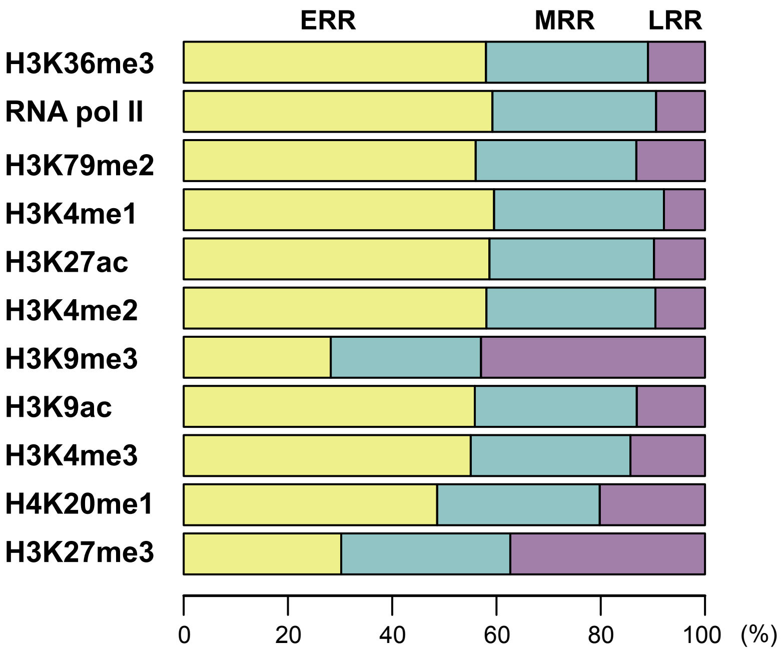


**Figure S5 Replication timing distribution of histone mark-enriched genomic regions**
For genomic bins with histone modification levels or RNA polymerase II binding above the median, the proportions of bins classified as ERR, MRR, and LRR are shown.

### Figure S6

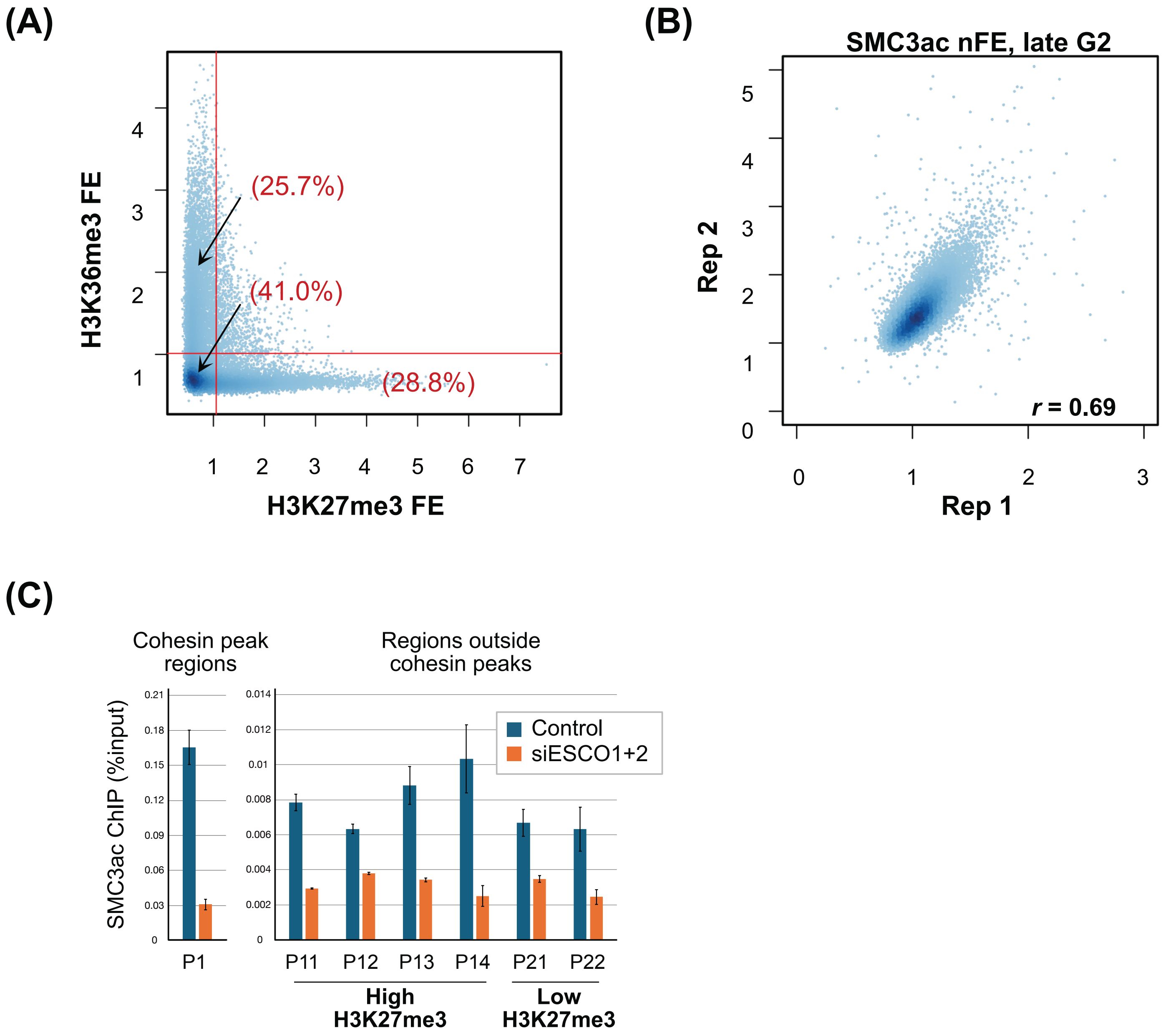


**Figure S6 Supplemental data supporting preferential retention of SMC3ac in H3K27me3-marked regions**

(A) Relationship between H3K27me3 and H3K36me3 levels across the genome. Each point represents a 100-kb genomic bin. Red lines indicate the genome-wide mean levels of H3K27me3 and H3K36me3. The percentages of the genome represented by regions with only H3K27me3 levels above the genome-wide mean, regions with only H3K36me3 levels above the genome-wide mean, and regions with both marks at or below their respective genome-wide means are indicated in parentheses. Together, these three categories account for 95.5% of the genome.

(B) Reproducibility of SMC3ac ChIP-seq profiles in late G2 cells. SMC3ac nFE values calculated in 100-kb bins were compared between two independent experiments. r, Pearson correlation coefficient.

(C) ChIP-qPCR analysis of chromosomal SMC3ac binding in late G2 cells. SMC3ac levels were measured in control cells and cells depleted of both ESCO1 and ESCO2 (siESCO1+2) at a cohesin peak (P1), non-peak regions with high H3K27me3 levels (P11–P14), and non-peak regions with low H3K27me3 levels (P21 and P22). Input and immunoprecipitated (IP) fractions were measured in duplicate by qPCR. Error bars indicate standard deviations calculated by error propagation.

### Figure S7

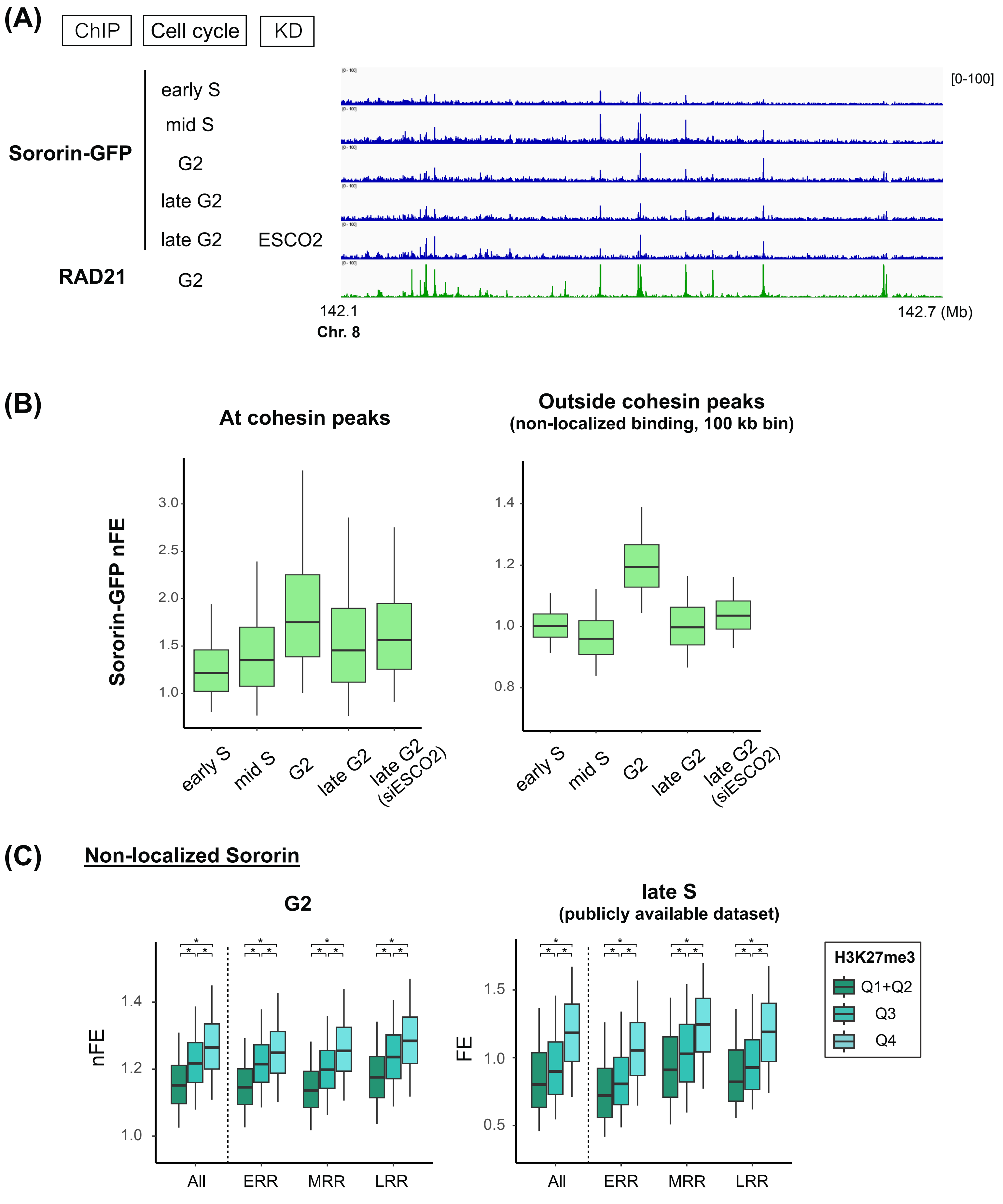

(A)
ChIP
Cell cycle
KD
Sororin-GFP
early S
mid S
G2
late G2
late G2
ESCO2
RAD21
G2
[0-100]
142.1
Chr. 8
142.7 (Mb)
(B)
At cohesin peaks
Outside cohesin peaks
(non-localized binding, 100 kb bin)
Sororin-GFP nFE
3.0
2.5
2.0
1.5
1.0
1.4
1.2
1.0
0.8
early S
mid S
G2
late G2
late G2 (siESCO2)
(C)
Non-localized Sororin
G2
late S
(publicly available dataset)
nFE
FE
1.4
1.2
1.0
1.5
1.0
0.5
All
ERR
MRR
LRR
H3K27me3
Q1+Q2
Q3
Q4

**Figure S7 Analysis of Sororin by calibrated ChIP-seq**

(A) Calibrated ChIP-seq profiles of Sororin-GFP. Read counts in the ChIP fractions at 100-bp resolution are shown, as in standard ChIP-seq analysis. Experimental conditions are indicated on the left. KD, knockdown by siRNA. RAD21 represents cohesin localization irrespective of its acetylation status.

(B) Quantification of chromosomal Sororin binding. ChIP-seq nFE values for cohesin peak-associated and non-localized Sororin are shown as boxplots. As for non-localized SMC3ac, non-localized Sororin nFE was calculated in 100-kb bins after excluding reads corresponding to cohesin peak regions. Boxes represent the interquartile range (25th–75th percentiles), whiskers indicate the 5th–95th percentiles, and horizontal lines within the boxes denote the median.

(C) Relationship between non-localized Sororin nFE and H3K27me3 levels. Based on H3K27me3 levels, the genome was divided into three groups (Q1+Q2: below the median, Q3: between the median and the third quartile, Q4: above the third quartile). Distributions of non-localized Sororin ChIP-seq nFE in G2 were compared across these groups. To examine whether this relationship was also observed in an independent dataset, previously reported Sororin ChIP-seq data from late S-phase cells (Ref. 41) were analyzed in the same manner (right panel). Boxes represent the interquartile range (25th–75th percentiles), whiskers indicate the 5th–95th percentiles, and bold lines denote the median. Data are shown for the whole genome (All) as well as for ERR, MRR, and LRR regions classified by replication timing. *, $P < 2.2 \times 10^{-16}$, two-sided Mann–Whitney *U* test.

#### Supplementary Table S1

Reagents and biological resources used in this study

| REAGENT or RESOURCE | SOURCE | IDENTIFIER |
|---|---|---|
| Antibodies | | |
| Mouse monoclonal anti-ESCO1 | Minamino *et al*. (2015) | N/A |
| Rabbit polyclonal anti-RAD21 | Minamino *et al*. (2015) | N/A |
| Mouse monoclonal anti-SMC3ac | Minamino *et al*. (2015) | N/A |
| Rabbit polyclonal anti-ESCO2 | Abcam | Cat# ab86003; RRID:AB_1924967 |
| Rabbit polyclonal anti-SMC1 | Abcam | Cat# ab21583; RRID:AB_2192477 |
| Rabbit monoclonal anti-SMC3 (clone D47B5) | Cell Signaling Technology | Cat# 5696, RRID:AB_10705575 |
| Rabbit monoclonal anti-MCM7 (clone D10A11) | Cell Signaling Technology | Cat# 3735, RRID:AB_2142705 |
| Mouse monoclonal anti-α-tubulin (clone B-5-1-2) | Sigma-Aldrich | Cat# T6074, RRID:AB_477582 |
| Rabbit monoclonal anti-Rpb1 (clone D8L4Y) | Cell Signaling Technology | Cat# 14958, RRID:AB_2687876 |
| Rabbit polyclonal anti-GFP | OriGene | Cat# TP401, RRID:AB_10013661 |
| Mouse monoclonal anti-H3K27me3 (clone MABI0323) | Fujifilm | Cat# 301-95253 RRID: AB_11123929 |
| Peroxidase-AffiniPure goat anti-Rabbit IgG | Jackson ImmunoResearch Labs | Cat# 111-035-144, RRID:AB_2307391 |
| Peroxidase-AffiniPure goat anti-Mouse IgG | Jackson ImmunoResearch Labs | Cat# 115-035-146, RRID:AB_2307392 |
| Chemicals, peptides, and recombinant proteins | | |
| D-MEM (High Glucose) | Fujifilm Wako | Cat# 044-29765 |
| Fetal Bovine Serum | Biosera | Cat# FB-1290/500 |
| Penicillin-Streptomycin-L-Glutamine Solution | Fujifilm Wako | Cat# 161-23201 |
| TrypLE Express Enzyme | Thermo Fisher Scientific | Cat# 12605028 |
| Thymidine | Sigma-Aldrich | Cat# T1895-10G |
| RO-3306 | Tocris | Cat# 4181 |
| EPZ-6438 | Selleck | Cat# S7128 |
| Lipofectamine RNAiMAX reagent | Thermo Fisher Scientific | Cat# 13778150 |
| cOmplete Protease Inhibitor Cocktail | Merck | Cat# 11697498001 |
| PhosSTOP | Merck | Cat# 4906845001 |
| Amersham Protran Premium 0.2 NC nitrocellulose Western blotting membrane | Cytiva | Cat# 10600004 |
| Luminata Forte Western HRP substrate | Merck | Cat# WBLUF0100 |
| Dynabeads Protein A | Thermo Fisher Scientific | Cat# DB10002 |
| Dynabeads Protein G | Thermo Fisher Scientific | Cat# DB10004 |
| RNase A | Merck | Cat# 10109142001 |

| | | |
|---|---|---|
| Proteinase K | Gold Biotechnology | Cat# P-480 |
| Critical commercial assays | | |
| QIAquick PCR Purification Kit | QIAGEN | Cat# 28106 |
| KAPA SYBR Fast qPCR Kit | KAPA Biosystems | Cat# KK4602 |
| NEBNext Ultra Ⅱ DNA Library Prep Kit for Illumina | New England Biolabs | Cat# E7645L |
| Anti-Mouse Detection Module | ProteinSimple | Cat# DM-002 |
| Anti-Rabbit NIR Detection Module | ProteinSimple | Cat# DM-007 |
| Cell lines | | |
| Human: HeLa S3 | JCRB Cell Bank | Cat# JCRB9010 |
| Mouse: C2C12 | ECACC | Cat# 91031101 |
| Human: HeLa expressing mouse Sororin with LAP tag | Nishiyama *et al*. (2013) | N/A |
| Model organisms | | |
| Mouse: R26R-H2B-EGFP | Abe *et al*. (2011) | CDB0203K |
| Mouse: FVB-Tg(Ddx4-cre)1Dcas/J (Vasa-Cre) | The Jackson Laboratory | RRID:IMSR_JAX:006954 |
| siRNA | | |
| siRNA target sequence: ESCO1: UGAAGUAUUUGUCUUUCAACACUGG | Thermo Fisher Scientific | Assay ID:HSS132872, Cat# 1299001 |
| siRNA target sequence: ESCO2: AUAACUUGCCAUCUGGUGUUGGGUC | Thermo Fisher Scientific | Assay ID:HSS152845, Cat# 1299001 |
| siRNA target sequence: MCM7: CGUCACUCGUGUCUCUGAAGUCAAA | Thermo Fisher Scientific | Assay ID:HSS106405, Cat# 1299001 |

**Supplementary Table S2**

Primer pairs used for qPCR
• Coordinates are based on hg19.
• Primer sequences are shown 5′ to 3′.

| Target locus ID | Chr | Start | End | Forward | Reverse |
|---|---|---|---|---|---|
| P1 | chr12 | 53278667 | 53278827 | TAGACGGAGCTGGAAGGAAA | AGGGACAGTCACCACCTTTG |
| P2 | chr17 | 32688528 | 32688678 | AAACCTCTGAGCTCATAATGCTG | CTGGGTGCAAGCCACACT |
| P3 | chr1 | 176529142 | 176529282 | GCAGTGTTGCTTCATTCACAGG | AAGGGATGCCAACGAACAAC |
| P4 | chr2 | 54931731 | 54931835 | TGTTGACATGTGCAGCTCAAG | TTGGATAGCTTGTTTGCTTTGC |
| P5 | chr3 | 119901853 | 119901972 | AATGCCCAGTGTCCCAAATG | TGGCGTCAGTTACCCAAAAG |
| P6 | chr12 | 40501367 | 40501558 | GCAAGGCTCTACCGTCATTC | CCTTCTCTTCAGAAGCCGTG |
| P7 | chr15 | 37426978 | 37427078 | TGTAGCAATGCTCTCTCACTCC | TGGGTCATCCACTTGGTATCAC |
| P8 | chrX | 32549072 | 32549144 | ACCCAAAAGCTGCCTATGTC | AGCTCTGAAAGGGAATGTGC |
| P11 | chr5 | 37854706 | 37854795 | CCCCTGTCACTGTATTGTGAAG | AGCAAGACACATGGCAAAGC |
| P12 | chr8 | 74029999 | 74030070 | TGTGCGCACTCACAGGCGAA | ACAGCCCAAAGCCTTGGCACC |
| P13 | chr6 | 110720591 | 110720670 | TGGAAGAGGCGTAGGTGGCAACA | CGACCCACATGGGACGCCAC |
| P14 | chr8 | 72568889 | 72569036 | CCCCACACAACTGTCTCTGTCACCT | ACACAACTGCAGCCACCGGA |
| P21 | chr11 | 94553607 | 94553702 | TTGCTGGCAGTGCTTCATAC | TGGTGCCAAGAAGGTTGGAG |
| P22 | chr6 | 38040873 | 38040959 | ATGGGAAAGGCTCAGCAAAGAGGC | GGCCACAGGTACTGTTGCGTGGA |

**Supplementary Table S3**

Summary of read mapping statistics

| Sample description | #Total reads | #uniquely mapped reads | | | | Occupancy Ratio | Normalization value used at the binning step | Relevant figures |
|---|---|---|---|---|---|---|---|---|
| | | To human (hg19) | | To mouse (mm10) | | | | |
| earlyS_ESCO2, ChIP | 27996629 | 23279782 | 83.15% | 3817457 | 13.64% | 1.2774 | 25547386 | Fig1AC, Fig2A-D, FigS2AE |
| earlyS_ESCO2, Input | 31481447 | 25223249 | 80.12% | 5283391 | 16.78% | NA | 20000000 | Fig1AC, Fig2A-D, FigS2AE |
| midS_ESCO2_rep1, ChIP | 38969551 | 33177654 | 85.14% | 4466626 | 11.46% | 1.2931 | 25862261 | Fig1AC, Fig2D, FigS2A-CE |
| midS_ESCO2_rep1, Input | 31711409 | 26212401 | 82.66% | 4563281 | 14.39% | NA | 20000000 | Fig1AC, Fig2D, FigS2A-CE |
| G2_ESCO2, ChIP | 31696139 | 26775804 | 84.48% | 3879427 | 12.24% | 0.8771 | 17542198 | Fig1AC, Fig2D, FigS2AE |
| G2_ESCO2, Input | 30553076 | 26250390 | 85.92% | 3335914 | 10.92% | NA | 20000000 | Fig1AC, Fig2D, FigS2AE |
| midS_ESCO2_rep2, ChIP | 36006885 | 30523784 | 84.77% | 4249227 | 11.80% | 1.5029 | 30058055 | FigS2C |
| midS_ESCO2_rep2, Input | 38778441 | 31099022 | 80.20% | 6506526 | 16.78% | NA | 20000000 | FigS2C |
| midS_siESCO2_ESCO2, ChIP | 36256926 | 28158080 | 77.66% | 7016431 | 19.35% | 0.8767 | 17533346 | Fig1A, FigS2C |
| midS_siESCO2_ESCO2, Input | 37394101 | 29774591 | 79.62% | 6504199 | 17.39% | NA | 20000000 | Fig1A, FigS2C |
| earlyS_MCM7_rep1, ChIP | 42735303 | 37214867 | 87.08% | 4013755 | 9.39% | 2.8974 | 57948797 | Fig1AC, Fig2ABD, FigS2BDE |
| earlyS_MCM7_rep1, Input | 45674833 | 33653207 | 73.68% | 10516598 | 23.02% | NA | 20000000 | Fig1AC, Fig2ABD, FigS2BDE |
| midS_MCM7, ChIP | 36001906 | 31298569 | 86.94% | 3405543 | 9.46% | 2.6241 | 52482258 | Fig1AC, Fig2D, FigS2E |
| midS_MCM7, Input | 40055798 | 30208377 | 75.42% | 8625252 | 21.53% | NA | 20000000 | Fig1AC, Fig2D, FigS2E |
| G2_MCM7, ChIP | 37824330 | 32565071 | 86.10% | 4027510 | 10.65% | 1.6381 | 32761318 | Fig1AC, Fig2D, FigS2E |
| G2_MCM7, Input | 36965855 | 29742911 | 80.46% | 6025588 | 16.30% | NA | 20000000 | Fig1AC, Fig2D, FigS2E |
| earlyS_MCM7_rep2, ChIP | 44295023 | 39446907 | 89.05% | 3303192 | 7.46% | 3.6745 | 73489234 | FigS2D |
| earlyS_MCM7_rep2, Input | 43454938 | 32152642 | 73.99% | 9893074 | 22.77% | NA | 20000000 | FigS2D |
| earlyS_siMCM7_MCM7, ChIP | 46967096 | 39932928 | 85.02% | 5368054 | 11.43% | 2.1367 | 42734314 | Fig1A, FigS2D |
| earlyS_siMCM7_MCM7, Input | 45323216 | 34002754 | 75.02% | 9766671 | 21.55% | NA | 20000000 | Fig1A, FigS2D |
| earlyS_RAD21, ChIP | 37558919 | 30452910 | 81.08% | 6071724 | 16.17% | 1.5653 | 31306104 | Fig4C |
| earlyS_SMC3ac, ChIP | 41366732 | 33323096 | 80.56% | 6383352 | 15.43% | 1.6292 | 32584330 | Fig3A-C, Fig4A, Fig5A, Fig6B, FigS4B |
| earlyS_RAD21/SMC3ac, Input | 35919675 | 26600446 | 74.06% | 8301779 | 23.11% | NA | 20000000 | Fig1B, Fig3A-C, Fig4A, Fig5A, Fig6B, FigS4B |
| earlyS_siESCO1_SMC3ac, ChIP | 40627452 | 31123950 | 76.61% | 7792496 | 19.18% | 1.2132 | 24263210 | Fig3C, Fig4A, Fig5A |
| earlyS_siESCO1_SMC3ac, Input | 38290301 | 28536989 | 74.53% | 8667788 | 22.64% | NA | 20000000 | Fig3C, Fig4A, Fig5A |
| earlyS_siESCO2_SMC3ac, ChIP | 40088867 | 30770312 | 76.76% | 7557174 | 18.85% | 1.3164 | 26327926 | Fig3C, Fig4A, Fig5A |
| earlyS_siESCO2_SMC3ac, Input | 36797528 | 27003723 | 73.38% | 8730474 | 23.73% | NA | 20000000 | Fig3C, Fig4A, Fig5A |
| earlyS_siESCO1&2_SMC3ac, ChIP | 40259257 | 27476464 | 68.25% | 10741597 | 26.68% | 0.8280 | 16560502 | Fig3C, Fig4A, Fig5A |
| earlyS_siESCO1&2_SMC3ac, Input | 38060254 | 27930385 | 73.38% | 9041249 | 23.76% | NA | 20000000 | Fig3C, Fig4A, Fig5A |
| midS_RAD21, ChIP | 38183615 | 30681683 | 80.35% | 6430558 | 16.84% | 1.2496 | 24991178 | Fig4C |
| midS_SMC3ac, ChIP | 38197108 | 32302958 | 84.57% | 4631611 | 12.13% | 1.8266 | 36531410 | Fig3A-C, Fig4A, Fig5A, Fig6B, FigS4B |
| midS_RAD21/SMC3ac, Input | 35133512 | 27095054 | 77.12% | 7096043 | 20.20% | NA | 20000000 | Fig1B, Fig3A-C, Fig4A, Fig5A, Fig6B, FigS4B |
| midS_siESCO1_SMC3ac, ChIP | 40278194 | 32734342 | 81.27% | 6253823 | 15.53% | 1.3454 | 26907125 | Fig3AC, Fig4A, Fig5A |
| midS_siESCO1_SMC3ac, Input | 38476897 | 29788846 | 77.42% | 7656547 | 19.90% | NA | 20000000 | Fig3AC, Fig4A, Fig5A |
| midS_siESCO2_SMC3ac, ChIP | 39916094 | 31964012 | 80.08% | 6495771 | 16.27% | 1.3343 | 26686149 | Fig3AC, Fig4A, Fig5A |
| midS_siESCO2_SMC3ac, Input | 36118713 | 27641998 | 76.53% | 7495399 | 20.75% | NA | 20000000 | Fig3AC, Fig4A, Fig5A |
| midS_siESCO1&2_SMC3ac, ChIP | 43998490 | 30796290 | 69.99% | 10452142 | 23.76% | 0.8659 | 17317016 | Fig3AC, Fig4A, Fig5A |
| midS_siESCO1&2_SMC3ac, Input | 36856543 | 27718462 | 75.21% | 8145525 | 22.10% | NA | 20000000 | Fig3AC, Fig4A, Fig5A |
| G2_RAD21, ChIP | 39002938 | 33266067 | 85.29% | 4569576 | 11.72% | 1.3517 | 27033226 | Fig3AB, Fig4C, FigS2AB |
| G2_SMC3ac, ChIP | 40540435 | 34757866 | 85.74% | 4338511 | 10.70% | 1.4875 | 29749846 | Fig3A-C, Fig4AB, Fig5A, Fig6AB, FigS2B, FigS4B |
| G2_RAD21/SMC3ac, Input | 38714248 | 31694280 | 81.87% | 5884685 | 15.20% | NA | 20000000 | Fig1B, Fig3A-C, Fig4A-C, Fig5A, Fig6AB, FigS2AB, FigS4B |
| G2_siESCO1_SMC3ac, ChIP | 40993679 | 34331438 | 83.75% | 4824845 | 11.77% | 1.4154 | 28307011 | Fig3C, Fig4AB, Fig5A |
| G2_siESCO1_SMC3ac, Input | 36494541 | 29546652 | 80.96% | 5877108 | 16.10% | NA | 20000000 | Fig3C, Fig4AB, Fig5A |
| G2_siESCO2_SMC3ac, ChIP | 39913920 | 32753298 | 82.06% | 5524340 | 13.84% | 1.2585 | 25170654 | Fig3C, Fig4AB, Fig5A |
| G2_siESCO2_SMC3ac, Input | 41022955 | 32870452 | 80.13% | 6977431 | 17.01% | NA | 20000000 | Fig3C, Fig4AB, Fig5A |
| G2_siESCO1&2_SMC3ac, ChIP | 40109527 | 31699433 | 79.03% | 6999559 | 17.45% | 0.9624 | 19247294 | Fig3C, Fig4AB, Fig5A |
| G2_siESCO1&2_SMC3ac, Input | 38903183 | 31149943 | 80.07% | 6619362 | 17.01% | NA | 20000000 | Fig3C, Fig4AB, Fig5A |
| lateG2_SMC3ac_rep1, ChIP | 40195549 | 33341356 | 82.95% | 5386531 | 13.40% | 0.8698 | 17396143 | Fig5A, Fig6A, FigS6B |
| lateG2_SMC3ac_rep1, Input | 35768332 | 30383357 | 84.94% | 4269575 | 11.94% | NA | 20000000 | Fig1B, Fig5A, Fig6A, FigS6B |
| G2_siRAD21_RAD21, ChIP | 41740686 | 15140785 | 36.27% | 25388196 | 60.82% | 0.2121 | 4241055 | Fig3B |
| G2_siRAD21_SMC3ac, ChIP | 35621020 | 22055338 | 61.92% | 12236141 | 34.35% | 0.6409 | 12818188 | Fig3B |
| G2_siRAD21_SMC3ac, Input | 40846457 | 31771424 | 77.78% | 7900311 | 19.34% | NA | 20000000 | Fig3B |
| Asy_Rpb1 (RNA pol II), ChIP | 37482563 | 30448540 | 81.23% | NA | NA | NA | 20000000 | Fig2AC, Fig6A |
| Asy_Rpb1 (RNA pol II), Input | 55793132 | 42466802 | 76.11% | NA | NA | NA | 20000000 | Fig2AC, Fig6A |
| lateG2_SMC3ac_rep2, ChIP | 74888861 | 60481288 | 80.76% | 7514676 | 10.03% | 1.8538 | 37076462 | Fig6BC, FigS6B |
| lateG2_H3K27me3, ChIP | 74392457 | 62399938 | 83.88% | 3458553 | 4.65% | 4.1557 | 83113875 | Fig6C |
| lateG2_SMC3ac(rep2)/H3K27me3, input | 81591847 | 59663855 | 73.12% | 13742423 | 16.84% | NA | 20000000 | Fig6BC, FigS6B |
| lateG2_EZP6438_SMC3ac_rep2, ChIP | 71,808,565 | 57,815,683 | 80.51% | 7,301,420 | 10.17% | 1.3981 | 27962569 | Fig6C |
| lateG2_EZP6438_H3K27me3, ChIP | 81,203,392 | 55,816,609 | 68.74% | 18,044,835 | 22.22% | 0.5462 | 10923410 | Fig6C |
| lateG2_EZP6438_SMC3ac(rep2)/H3K27me3, input | 70,432,024 | 53,799,358 | 76.38% | 9,499,080 | 13.49% | NA | 20000000 | Fig6C |
| earlyS_Sororin-GFP, ChIP | 257208002 | 107289812 | 41.71% | 140044597 | 54.45% | 0.1962 | 3923736 | FigS7A-C |
| earlyS_Sororin-GFP, input | 65902510 | 48145718 | 73.06% | 12330138 | 18.71% | NA | 20000000 | FigS7A-C |
| midS_Sororin-GFP, ChIP | 75417894 | 33438811 | 44.34% | 38829101 | 51.49% | 0.2073 | 4146310 | FigS7A-C |
| midS_Sororin-GFP, input | 88458437 | 65141024 | 73.64% | 15682387 | 17.73% | NA | 20000000 | FigS7A-C |
| G2_Sororin-GFP, ChIP | 77564486 | 42001076 | 54.15% | 30403831 | 39.20% | 0.237 | 4739944 | FigS7A-C |
| G2_Sororin-GFP, input | 73833738 | 56769441 | 76.89% | 9739014 | 13.19% | NA | 20000000 | FigS7A-C |
| lateG2_Sororin-GFP, ChIP | 62910605 | 29620140 | 47.08% | 30350384 | 48.24% | 0.1972 | 3943158 | FigS7A-C |
| lateG2_Sororin-GFP, input | 76196817 | 57444414 | 75.39% | 11604561 | 15.23% | NA | 20000000 | FigS7A-C |
| lateG2_siESCO2_Sororin-GFP, ChIP | 81691563 | 37090626 | 45.40% | 41069491 | 50.27% | 0.2102 | 4204611 | FigS7A-C |
| lateG2_siESCO2_Sororin-GFP, input | 130870222 | 96312850 | 73.59% | 22421014 | 17.13% | NA | 20000000 | FigS7A-C |

#### Supplementary Table S4

**Publicly available sequencing data used in this study**

All datasets were derived from HeLa S3 cells.

| Description | File ID/ Accession ID | Download site |
|---|---|---|
| GFP-ESCO1, ChIP-seq in G2 | SRR1916173 | https://trace.ncbi.nlm.nih.gov/Traces?run=SRR1916173 |
| Replication timing by Repli-seq (wavelet-smoothed signal) | GSM923449_hg19_wgEncodeUwRepliSeqHelas3WaveSignalRep1.bigWig | https://www.ncbi.nlm.nih.gov/geo/download/?acc=GSM923449&format=file&file=GSM923449%5Fhg19%5FwgEncodeUwRepliSeqHelas3WaveSignalRep1%2EbigWig |
| H3K4me1, ChIP-seq | E117-H3K4me1.tagAlign.gz | https://egg2.wustl.edu/roadmap/data/byFileType/alignments/consolidated/ |
| H3K4me2, ChIP-seq | E117-H3K4me2.tagAlign.gz | |
| H3K4me3, ChIP-seq | E117-H3K4me3.tagAlign.gz | |
| H3K9ac, ChIP-seq | E117-H3K9ac.tagAlign.gz | |
| H3K27ac, ChIP-seq | E117-H3K27ac.tagAlign.gz | |
| H3K27me3, ChIP-seq | E117-H3K27me3.tagAlign.gz | |
| H3K36me3, ChIP-seq | E117-H3K36me3.tagAlign.gz | |
| H3K79me2, ChIP-seq | E117-H3K79me2.tagAlign.gz | |
| H4K20me1, ChIP-seq | E117-H4K20me1.tagAlign.gz | |
| Input for ChIP (roadmap) | E117-Input.tagAlign.gz | |
| H3K9me3, ChIP-seq (Fold change over control) | ENCFF761QZP | https://www.encodeproject.org/files/ENCFF761QZP/ |
| Sororin, ChIP-seq in late S | ERR1194877 | https://www.ebi.ac.uk/ena/browser/view/ERR1194877 |
| Genome control for Sororin ChIP | ERR1195873 | https://www.ebi.ac.uk/ena/browser/view/ERR1195873 |
| RNA-seq | SRR16555324 | https://trace.ncbi.nlm.nih.gov/Traces/?view=run_browser&acc=SRR16555324&display=download |
